# A systematic targeted genetic screen identifies proteins involved in cytoadherence of the malaria parasite *P. falciparum*

**DOI:** 10.1101/2024.07.24.604743

**Authors:** Nina Küster, Lena Roling, Ardin Ouayoue, Katharina Steeg, Jude M Przyborski

**Affiliations:** University of Giessen

**Keywords:** *Plasmodium falciparum*, red blood cells, RBC, knobs, malaria, virulence, *Pf*EMP1, cytoadherence

## Abstract

Immediately after invading their chosen host cell, the mature human erythrocyte, malaria parasites begin to export an array of proteins to this compartment, where they initiate processes that are prerequisite for parasite survival and propagation, including nutrient import and immune evasion. One consequence of these activities is the emergence of novel adhesive phenotypes that can lead directly to pathology in the human host. To identify parasite proteins involved in this process we used modern genetic tools to target genes encoding 15 exported parasite proteins, selected by an *in-silico* workflow. This resulted in 4 genetically modified parasite lines that were then characterised in detail. Of these lines, 3 could be shown to have aberrations in adhesion, and of these 1 appears to have a block in the transport and/or correct folding of the major surface adhesin PfEMP1 (*Plasmodium falciparum* erythrocyte membrane protein 1). Our data expand the known factors involved in this important process, and once again highlight the complexity of this phenomenon.

## 1. INTRODUCTION

*Plasmodium falciparum* is the primary cause of the most severe form of malaria, known as malaria tropica. It accounts for >200 million clinical cases and >600,000 deaths annually, predominantly affecting children under the age of 5 in sub-Saharan Africa (WHO, 2023). The severe pathology of malaria is primarily due to the infected red blood cells (iRBCs) adhering to various receptors on endothelial cells in tissues and organs (Miller et al., 2002). This cytoadherence leads to reduced blood flow, hypoxia, and, in cases of cerebral malaria, increased intracranial pressure (Miller et al., 2002; Waller et al., 1991; Newton et al., 1991). Cytoadherence is a result of parasite-induced modifications in host cells, where parasite-encoded proteins are transported to the surface of the iRBCs, facilitating endothelial binding and antigenic variation, with the major mediator in this process being the *Plasmodium falciparum* erythrocyte membrane protein 1 (PfEMP1) (Su et al., 1995; Baruch et al., 1995; Smith et al., 1995). Besides these surface proteins, *P. falciparum* also encodes, expresses and exports numerous other proteins to the iRBCs, a number of which underpin cytoadherence through molecular mechanisms that are still poorly understood (Maier et al., 2008; Maier et al., 2009; Sargeant et al., 2006). The function and importance of further exported proteins remains unknown, in part due to limitations in techniques for genetic manipulation of the parasite. In this current study, we utilised several recently developed genetic tools to address this and identify several exported proteins of previously unknown function which play a role, likely indirectly, in cytoadherence (Birnbaum et al., 2017; Prommana et al., 2013). Our data further expand knowledge of the factors required for cytoadherence and highlight once again the large number of ancillary parasite proteins required for this essential process. Furthermore, our carefully controlled multi-technique strategy identifies a number of genes which appear, within the limitations of the genetic tools used, to be essential for blood stage parasite survival.

## 2. RESULTS

### 2.1 Selection of genes for analysis

We utilised an *in-silico* workflow to identify genes likely involved in host-cell modification, and which could be analysed using the selection-linked integration technique (SLI) method described by Birnbaum and colleagues (Birnbaum et al., 2017). Our criteria for inclusion (based on data available at PlasmoDB [Alvarez-Jarreta et al., 2024]) were 1. gene product predicted to be exported 2. not part of a large multi-gene family (thus excluding *rif*, *stevor*, *var*, *phist* and *fikk* genes) 3. not previously knocked out 4. evidence for high transcript levels at 0-16 and 44-48 hours post invasion (hpi) 5. coding sequence >600 bp. This strategy resulted in a list of 15 genes which were then carried on to further analysis (Table 1). Each of these genes (as expected) encodes a protein containing a putative PEXEL export sequence (Supplementary 1), and PF^0300^ and PF^0600^ are predicted to contain 2, respectively 1 membrane crossing domain(s) (Supplementary 1). The two larger proteins PF^1600^ and PF^3300^ contain degenerate tandem repeats in the central and c-terminal region, a common feature in exported parasite proteins but whose significance has not yet been determined. Alignments of homologues from all *P. falciparum* strains deposited at PlasmoDB reveal a high level of conservation amongst all strains, however the number of c-terminal repeats appears to be flexible (Supplementary 1).

**TABLE 1.** Overview of GOI, genetic manipulation approaches and outcomes.

### 2.2 Generation of genetically modified parasite lines

We initially targeted all selected genes for disruption using the SLI-targeted gene disruption method (SLI-TGD, Supplementary 2 [Birnbaum et al., 2017]). This method utilises a two-step drug selection and only parasites with integration into the correct gene locus should result. As it appeared likely that some genes would be essential and thus resistant to this technique, we also utilised SLI to introduce a GFP-tag followed by a glmS ribozyme (Prommana et al., 2013) to enable us to manipulate protein levels. As GFP is a large tag and can interfere with protein function, several genes which were refractive to the introduction of GFP were tagged with a 3xHA-tag. As parental clone, we chose CS2, which can be selected to express a specific PfEMP1 variant (PfEMP1^var2csa^ [Reeder et al., 1999]), facilitating phenotypic analysis. A significant number of transgenic parasite lines were generated carrying episomal plasmids (Table 1). Following second selection, clonal integrant lines were generated using limiting dilution, and PCR analysis was used to verify integration into the correct gene locus (Supplementary 2, Supplementary 3). This revealed that, in addition to correct integration events, many parasite lines still retained episomal plasmids, which however are not expected to interfere with following experiments. Using this strategy, we were able to manipulate 2 genes to introduce GFP_glmS, and a further 6 loci could be obtained with an integrated 3xHA*_*glmS (Table 1). As a suitable control for the glmS lines obtained, we also generated parasites which had integrated a sequence which would generate a non-functional ribozyme, referred to as M9 (Prommana et al., 2013). In total, we generated 47 episomal and 16 integrant parasite lines (Table 1, M9 lines not listed). Excluding duplications obtained using multiple methods, and 2 genes whose analyses were published during the course of our study, we were left with 4 integrant parasite lines (and M9 controls) for further study. We refer to these genes/gene products using a short form of their PlasmoDB accession number for easy comprehension (Table 1, right hand column). As exported proteins are less likely to be essential than those involved in the parasite’s core metabolism (Maier et al., 2008), we were surprised that we were not able to inactivate any gene using the SLI-TGD method. This method has been shown to be highly dependable and (Birnbaum et al., 2017), in our hands, has even enabled inactivation of a gene (*pfa66*) previously thought essential (Diehl et al., 2021). Nevertheless, to control for potential technical reasons for this negative result, we re-cloned the gene targeting segment for *pfa66* we had originally used (Diehl et al., 2021) into the backbone of the vector used in this current analysis. This controls for potential rearrangements in the plasmid backbone which may not be immediately recognised despite detailed restriction analysis. We were able to obtain parasites which integrated this vector into the correct *pfa66* locus, thus verifying the validity of the experimental approach and technology, and fidelity of the vector used (data not shown). It appears that we may have fortuitously selected only genes which appear, according to this analysis, to be essential for parasite survival in blood stage culture.

### 2.3 Localisation of the gene products

As our strategy included incorporating a c-terminal tag into the final expressed protein, we could use this tag to localise the gene products. We used immunofluorescence assays (IFAs) to visualise the distribution of the 3xHA-tagged protein of interest (POI) in erythrocytes infected with our transgenic parasite lines (Figure 1A). To allow visualisation in 3 dimensions, we utilised a fixation protocol which preserved cell morphology, and collected Z-stacks (which are presented here as 2D maximal projection Z-project images). PF^0300^ revealed a “dotted” fluorescent pattern in the host cell (Figure 1A, upper panel). For protein PF^0600^, we could verify export to the host cell and observed an even distribution across the entire host cell (Figure 1A, middle panel). Of note, the fluorescence was also present in the area of the parasite body, highly indicative of a localisation below the erythrocyte plasma membrane (rather than the typical pattern of a soluble reporter in the host cell, which shows lack of or reduced fluorescence in this area). Our interpretation is supported by presentation of a single slice from the Z-series, collected from the middle of the infected cell, which shows a characteristic rim of fluorescence following the contours of the host cell plasma membrane, and a movie showing the entire Z-stack (Supplementary 4, Supplementary 5). PF^1600^ was exported to the host cell, and also located to fluorescent foci within the host cell (Figure 1A, lower panel). Despite numerous attempts to localise PF^3300^, we were not able to detect any specific signal, likely due to low expression levels. This phenomenon has previously been reported when relying on expression from an endogenous promoter (Birnbaum et al., 2017). The localisation of PF^0300^ and PF^1600^ was indicative of Maurer’s clefts (MC), parasite-induced structures in the infected host cell, which contain many exported proteins (Przyborski et al., 2005; Wickham et al., 2001; Hanssen et al., 2008; Lanzer et al., 2006; Przyborski et al., 2003). We therefore carried out co-labelling analyses using antisera against the MC resident protein PfSBP1 (Figure 1B). In neither case (Figure 1B, upper and lower panels) did we see significant overlap, or similar pattern of signals from the tagged POI and PfSBP1, making it unlikely that these fluorescent structures represent the MC.

**FIGURE 1.**
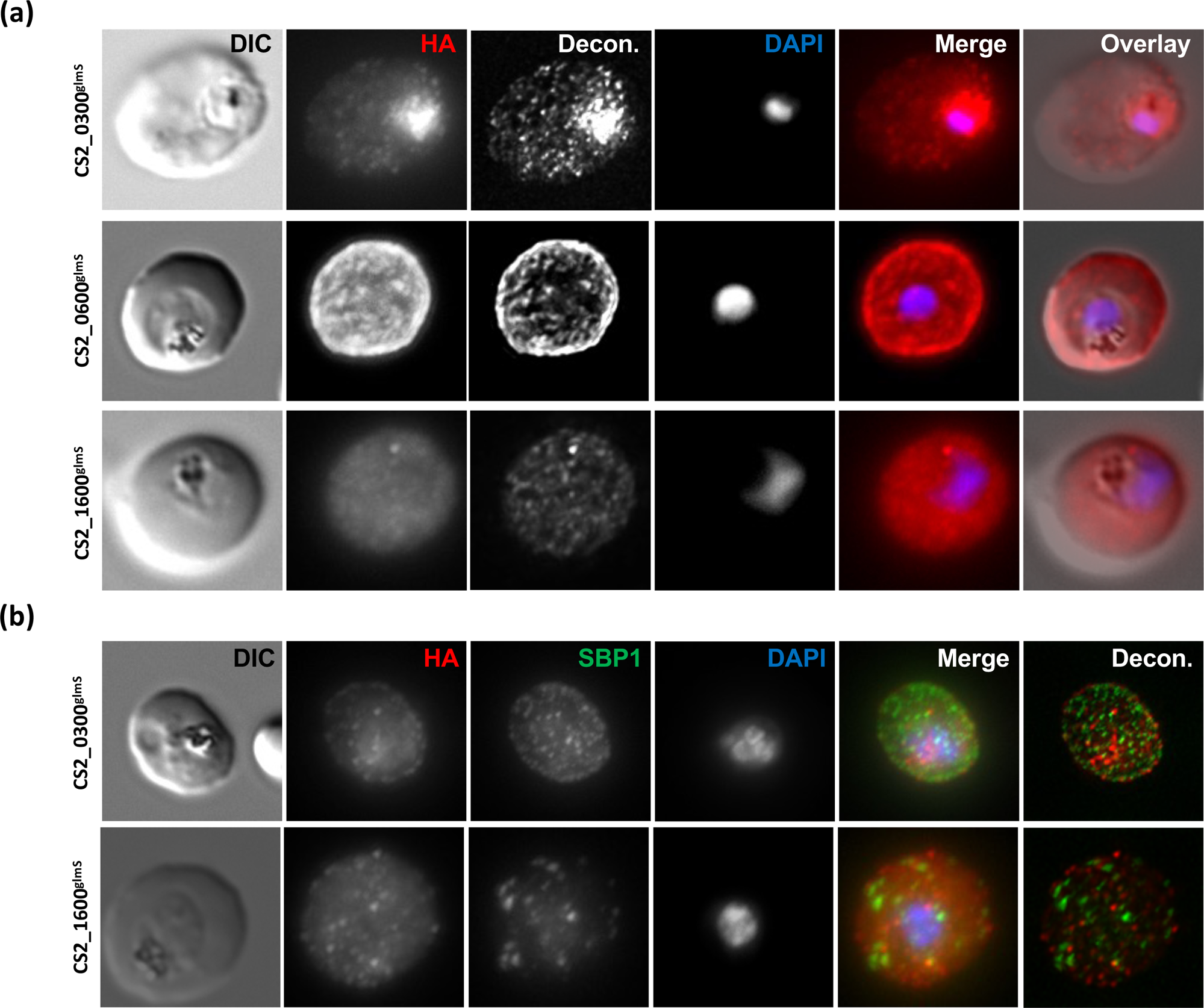
Localisation of POI. Fluorescence channels are shown individually in black/white for highest contrast. All images are maximal projections of Z-stack serial sections. Nuclear DNA was stained with DAPI. All images are representative of at least 20 independent observations. (a) Immunofluorescent localisation of POI. Fixed cells were treated with anti-HA antibodies to detect the POI, followed by suitable secondary antibodies. DIC, differential interference contrast; HA, anti-HA; Decon., maximal projection of deconvoluted Z-stack images; DAPI, nucleus. (b) Co-immunofluorescence analysis. Fixed cells were treated with anti-HA antibodies to detect the POI and anti-PfSBP1 antibodies followed by suitable secondary antibodies. DIC, differential interference contrast; HA, anti-HA; SBP1, anti-PfSBP1; DAPI, nucleus; Decon., maximal projection of deconvoluted Z-stack images.

### 2.4 Addition of glucosamine (GlcN) leads to a specific reduction in abundance of the POI

We wished to verify if incorporation of the glmS ribozyme into the endogenous gene locus would allow us to reduce abundance of the POI. We thus carried out western blot analysis using anti-HA antibodies. In parallel, we used parasite lines containing a non-functional ribozyme (M9) to control for the reported potentially toxic effect of GlcN and used parasite aldolase as a loading control for quantification. In the CS2_0300^glmS^ cell line, we observed a significant reduction in protein abundance compared to the control at concentrations ≥ 2.5 mM, which reduced protein abundance by 64.5+/−7.2%. Addition of higher concentrations of GlcN did not lead to any further dramatic drop in protein abundance. Parasites expressed a protein at the predicted molecular mass, and we could not detect any residual unskipped POI-NEO (Figure 2A). In the CS2_0600^glmS^ cell line, already addition of 1.25 mM GlcN led to a dramatic reduction (60.9+/−10.6%) in protein abundance compared to the paired control line, and parasites expressed the protein at the expected molecular mass, with minimal amounts of unskipped POI-NEO fusion detectable. Use of higher GlcN concentrations further decreased protein abundance (Figure 2B). In the CS2_1600^glmS^ cell line, we were able to show a significant reduction of protein abundance at GlcN concentrations ≥ 1.25 mM (58.3+/−10.3%), with a low abundance of unskipped POI-NEO also being detected. Unexpectedly, the POI showed an aberrant motility, running at <100 kDa compared to the calculated molecular mass of ≈80 kDa (Figure 2C). This phenomenon may be explained by the unusually low predicted pI of this protein (4.77), or potentially by uncharacterised stable posttranslational modification. In the CS2_3300^glmS^ cell line, addition of GlcN led to a definite reduction in protein abundance, however (and in accordance with our previous IFA results), the low level of protein expression even without the addition of GlcN did now allow for reliable quantification. Additionally, we note a not insignificant level of POI-NEO present in this cell line, indicative of suboptimal ribosomal skipping (Figure 2D).

**FIGURE 2.**
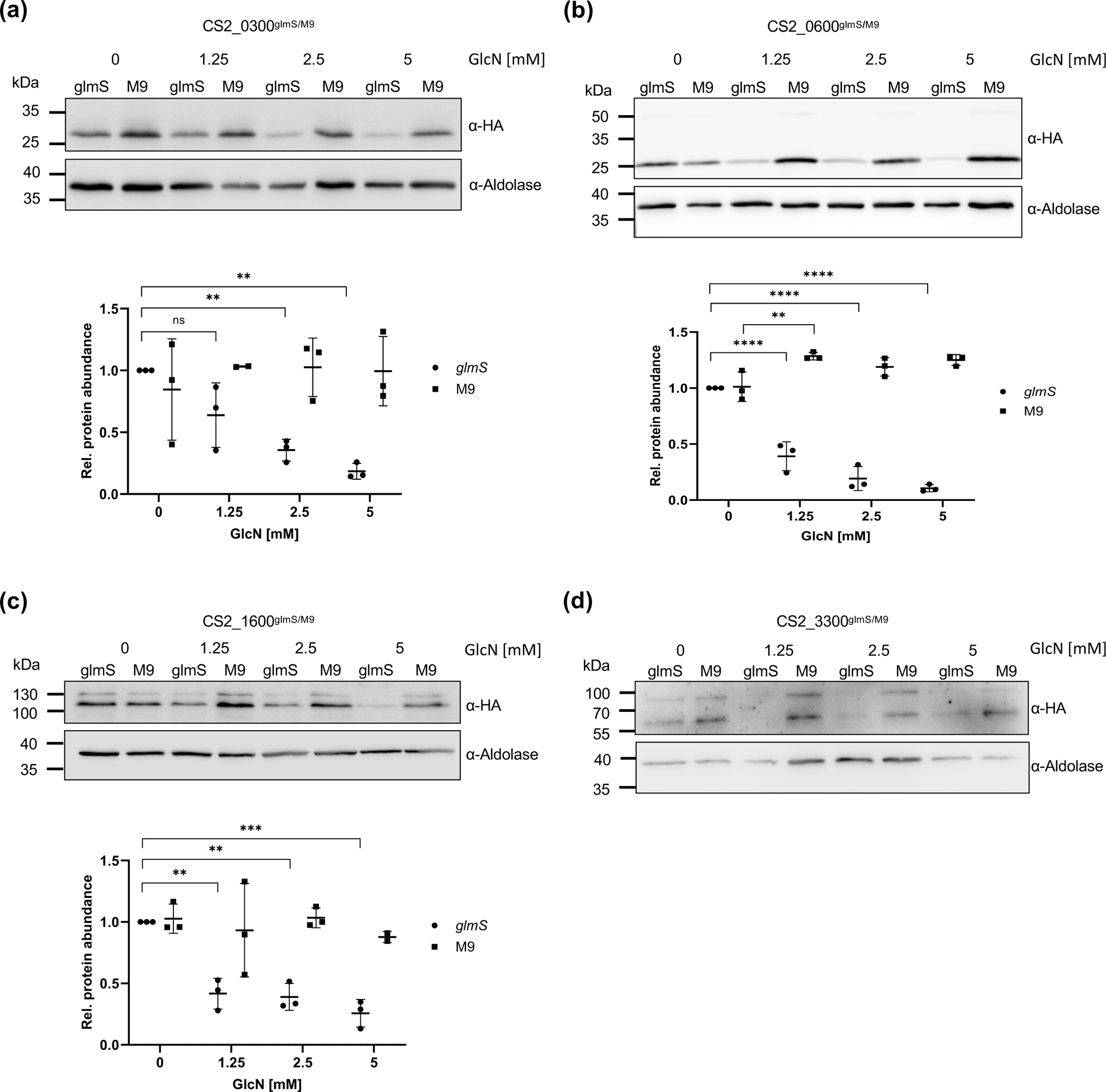
Effect of GlcN treatment on abundance of POI. Immunoblot detection of HA-tagged POI. GlmS and M9 cell lines were treated for 72 h with the GlcN concentration indicated. Anti-aldolase was used as a loading control. Protein abundance was quantified using ImageJ and statistically analysed with GraphPad Prism. N=3; ns, not significant; ** p<0.01; *** p<0.001; **** p<0.0001 (Student’s t-test).

### 2.5 Reduction in POI abundance does not lead to a reduction in parasite proliferation, life cycle progression or merozoite numbers

We analysed parasite proliferation in cell culture using the SybrGreen assay (Bennett et al., 2004; Smilkstein et al., 2004). As previously, we used parasite lines containing a non-functional ribozyme (M9) to control for the toxic effect of high levels of GlcN. Under these conditions, no parasite lines were shown to have a significantly lower rate of growth over 72 h when compared to the paired controls (Figure 3A-D). There was no significant difference in merozoite numbers (Figure 3A-D), in parasite morphology or stage progression as determined by examination of Giemsa-stained smears (Supplementary 6). We did however detect a (glmS-independent) effect of even 1.25 mM GlcN on parasite growth. In 3 of the 4 parasite lines (and paired controls), we noted significant reduction in growth compared to the non-GlcN treated controls (Figure 3A, B, D). When increasing the concentration to 2.5 mM, all studied cell lines showed a significant growth defect when compared to controls. This data highlights again the significance of robust experimental controls (i.e. M9) when using the glmS system, as non-specific effects may lead to incorrect data interpretation.

**FIGURE 3.**
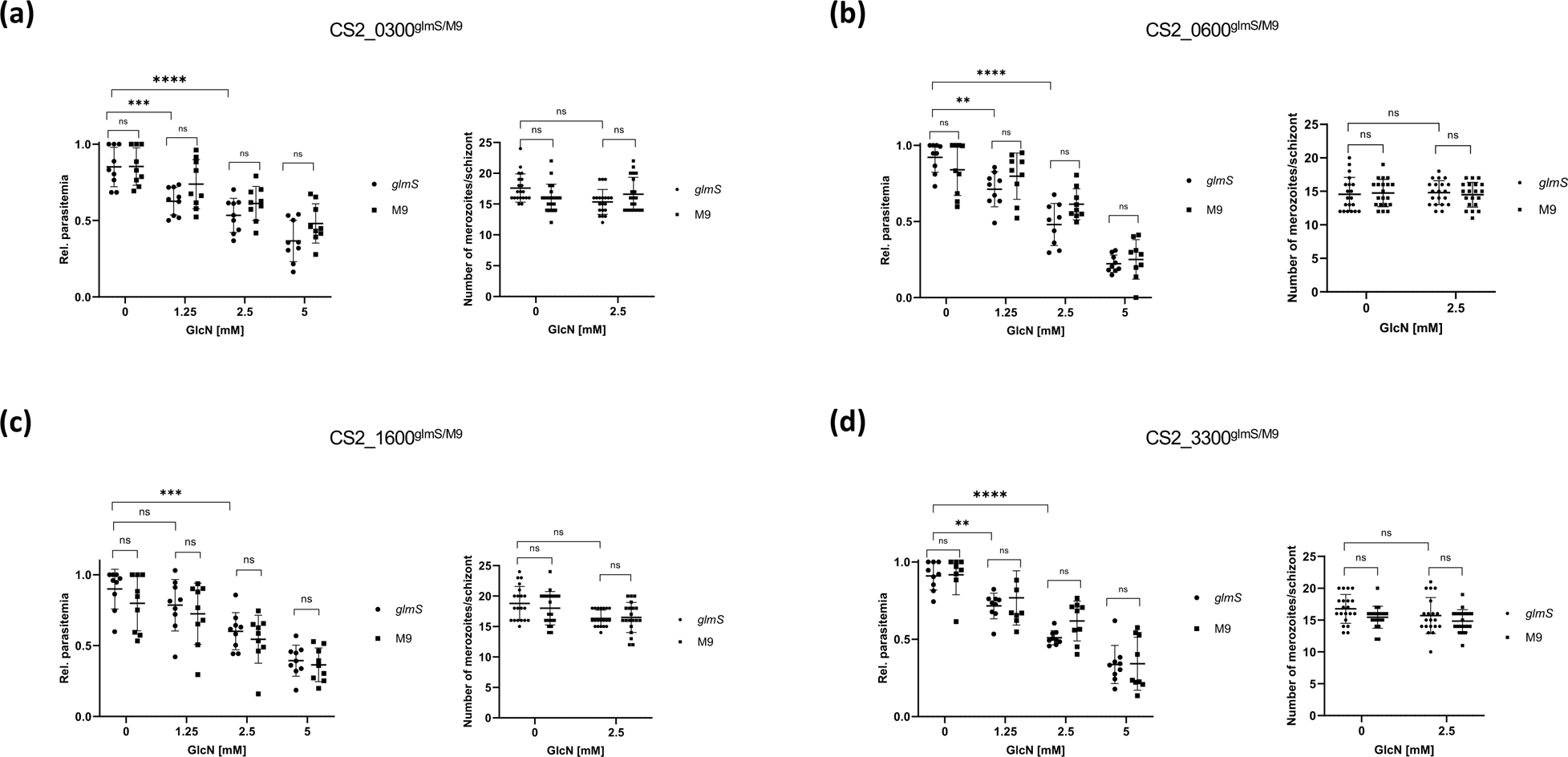
Effect of GlcN treatment on parasite proliferation and merozoite numbers. GlmS and M9 cell lines were treated for 72 h with the GlcN concentration indicated. Parasite proliferation was quantified using SybrGreen and merozoite numbers were quantified by visual examination of Giemsa-stained slides. Data were statistically analysed with GraphPad Prism. N=3; ns, not significant; ** p<0.01; *** p<0.001; **** p<0.0001 (Student’s t-test).

### 2.6 Reduction in POI abundance does not lead to specific defects in protein export

Several groups have highlighted that, even if exported proteins are not essential for parasite survival, their inactivation can have significant effects on other processes such as protein export or correct localisation of further exported proteins (Maier et al., 2008; Maier et al., 2009; Diehl et al., 2021; McHugh et al., 2020). For this reason, we studied the localisation of 4 further exported proteins by immunofluorescence. GlmS lines were treated with 2.5 mM GlcN for 72 h, fixed and probed with antibodies against PfKAHRP (localised to the knobs, a kind gift of Paul Gilson), PfSBP1 (localised to the MC, a kind gift of Catherine Braun-Breton), REX1 (MC, a kind gift of Matt Dixon) and PfEMP3 (localised to the inner leaflet of erythrocyte plasma membrane, a kind gift of Alan Cowman). Accounting for normal slight differences between individual infected cells, GlcN treatment did not lead to any incorrect localisation of the marker proteins (Figure 4, M9 controls shown in Supplementary 7). We cannot formally exclude that the localisation of other exported proteins would be disrupted upon GlcN treatment, but we think our results show that the most important protein export pathways still function with high fidelity.

**FIGURE 4.**
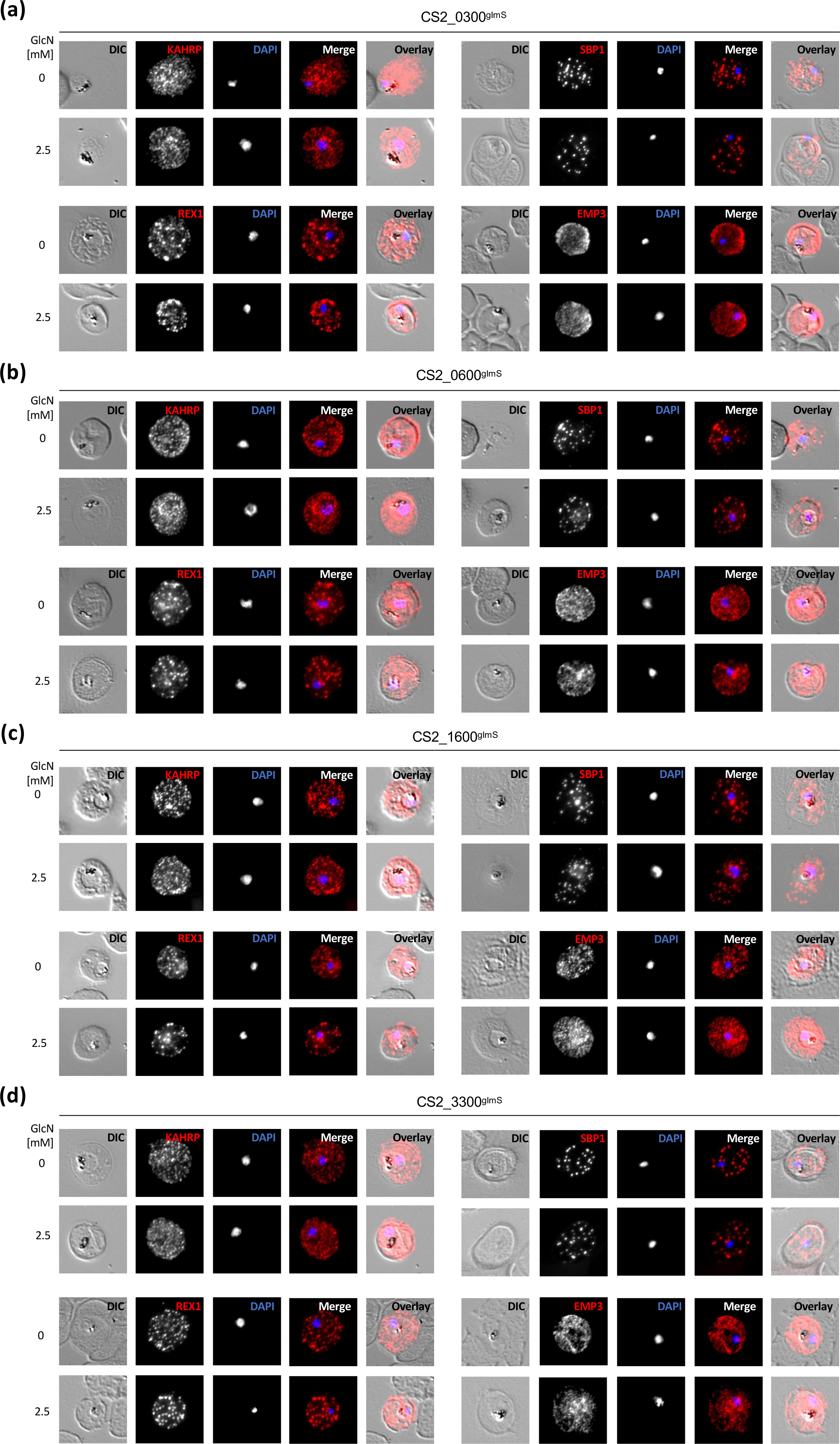
Localisation of other exported proteins. Immunofluorescent localisation of PfKAHRP, PfSBP1, PfREX3 and PfEMP3 in glmS cell lines +/− 2.5 mM GlcN treatment for 72 h. Cells were fixed and incubated with anti-PfKAHRP, anti-PfSBP1, anti-PfREX3 or anti-PfEMP3 respectively followed by Cy3-conjugated secondary antibody. Fluorescence channels are shown individually in black/white for highest contrast. DIC, differential interference contrast; DAPI, nucleus. All images are representative of at least 20 independent observations.

### 2.7 Transmission electron microscopy reveals no striking abnormality in host cell modification upon GlcN treatment

Upon entering human erythrocytes, malaria parasites modify the cellular architecture of the infected cell and induce the generation of several novel structures including the aforementioned Maurer’s clefts, and electron-dense protrusion of the erythrocyte plasma membrane referred to as knobs (Maier et al., 2008). To visualise any defects in the biogenesis of these structures, we treated parasites with GlcN as before, parasites were then fixed and prepared for transmission electron microscopy (TEM). As before, we analysed the M9 and glmS lines in parallel. In all the infected erythrocytes we studied, Maurer’s clefts (black inset) appeared to have a normal, elongated lamella morphology and did not show any evidence of abnormal vesiculation. Their positioning at the periphery of the infected cell orientated along the axis of the erythrocyte plasma membrane also appeared to be only subject to the expected natural variation (Figure 5A-D). Although we did not quantify knob structures, they could also be seen in all samples (red inset) and appeared structurally similar. We could see no evidence of deformed or extended knobs (Figure 5A-D).

**FIGURE 5.**
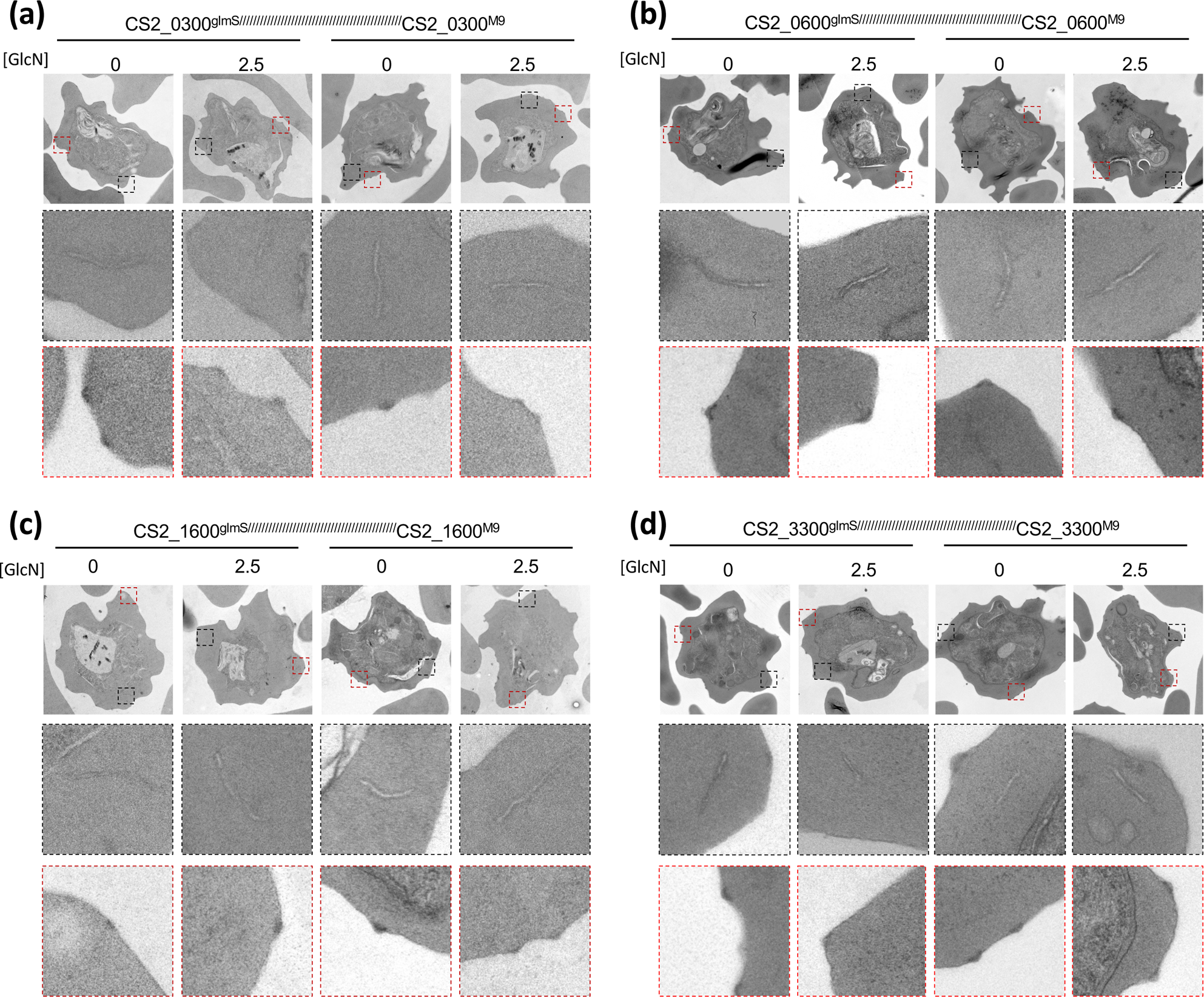
Transmission electron microscopy of parasite-infected erythrocytes. Erythrocytes infected with glmS and M9 cell lines, +/− 2.5 mM GlcN treatment for 72 h, were imaged and visualised using transmission electron microscopy. Black inset, magnified view of Maurer’s clefts morphology; Red inset, magnified view of knob morphology. All images are representative of at least 10 independent observations.

### 2.8 PF^0300^, PF^0600^ and PF^1600^ underpin cytoadherence

As has been previously demonstrated, many exported proteins are involved in host cell modifications that lead to the phenomenon of cytoadherence: the binding of infected erythrocytes to specific receptors on endothelial cells (Maier et al., 2008). Cytoadherence underlies how parasites cause pathology in the human host and thus is an important phenotype to study when generating mutant parasite lines. For our genetic analysis, we deliberately chose a parasite strain (CS2) which could be selected to express a specific PfEMP1 variant (PfEMP1^var2csa^) that is known to bind to chondroitin-4-sulphate (CSA), and this can be quantified by an *in vitro* binding assay (Reeder et al., 1999). All parasite lines used for these analyses were freshly selected for CSA binding prior to setting up the assay to ensure continued expression of PfEMP1^var2csa^. CSA was immobilised on a Petri dish, infected erythrocytes allowed to bind, and washing steps removed unbound infected erythrocytes. As a control for the specificity of binding, we included a control in which infected erythrocytes were allowed to bind in the presence of soluble CSA (CSA SOL). This analysis revealed that 3 of the 4 glmS strains studied showed, upon addition of GlcN, significant differences in CSA binding when compared to the untreated control (Figure 6A-C). CS2_0300^glmS^, CS2_0600^glmS^ and CS2_1600^glmS^ lines showed a reduction in binding of respectively 60.3+/−17.9%, 50.2+/− 34.6% and 45.3+/−20.3%. The CS2_3300^glmS^ line appears to show slightly reduced binding, but this was not statistically significant (Figure 6D).

**FIGURE 6.**
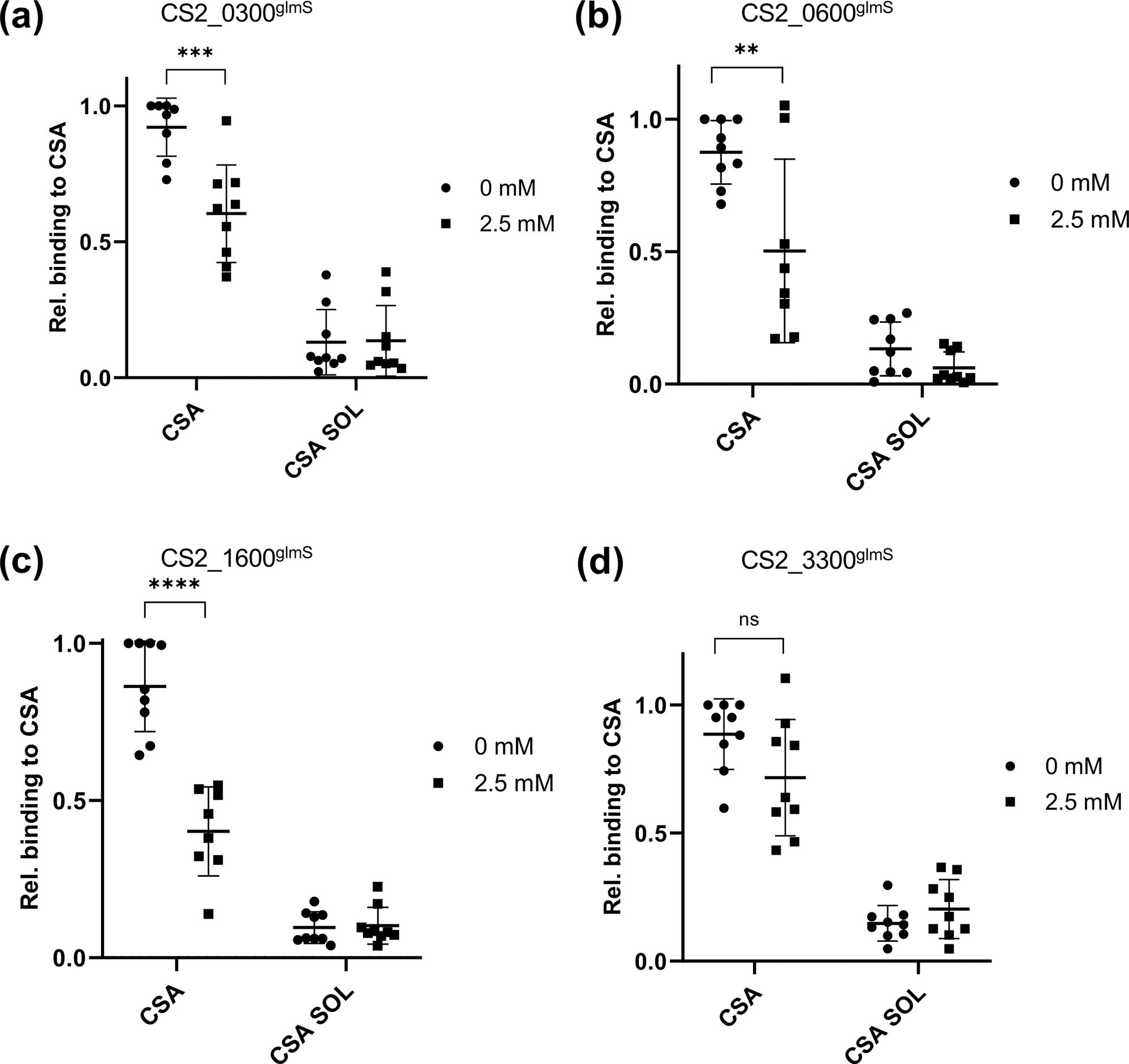
Static binding assay of parasite-infected erythrocytes. Erythrocytes infected with glmS and M9 cell lines, +/− 2.5 mM GlcN treatment for 72 h, were analysed for their ability to bind to immobilized CSA receptor. Captured images of Giemsa-stained cells were evaluated using Illastik, and the results statistically analysed with GraphPad Prism. N=3; ns, not significant; ** p<0.01; *** p<0.001; **** p<0.0001 (Student’s t-test).

### 2.9 Erythrocytes infected with CS2_0300^glmS^ parasite line show reduced surface exposure of PfEMP1^var2csa^

There are several things that could lead to a reduction in cytoadherence: i) reduced levels of PfEMP1 on the cell surface, possibly via blockage of its trafficking pathway ii) aberrant knob morphology leading to incorrect PfEMP1 presentation or iii) incorrect folding of PfEMP1. As we had demonstrated that all our parasite lines appeared to have normal Maurer’s cleft and knob morphology, we were interested to investigate if the reduced cytoadherence we observed could be explained by a reduction of PfEMP1^var2csa^ at the surface of the infected erythrocyte. Erythrocytes infected with synchronised trophozoite-stage parasites were treated with specific anti-var2CSA antibodies (P11, a kind gift of Benoit Gamain) followed by fluorescently conjugated secondary antibodies and relative surface fluorescence measured using flow cytometry. Erythrocytes infected with CS2_1600^glmS^ showed no significant decrease in surface fluorescence when treated with GlcN (Figure 7B), however erythrocytes infected with CS2_0300^glmS^ (Figure 7A) showed a highly significant reduction in surface detection of PfEMP1^var2csa^.

**FIGURE 7.**
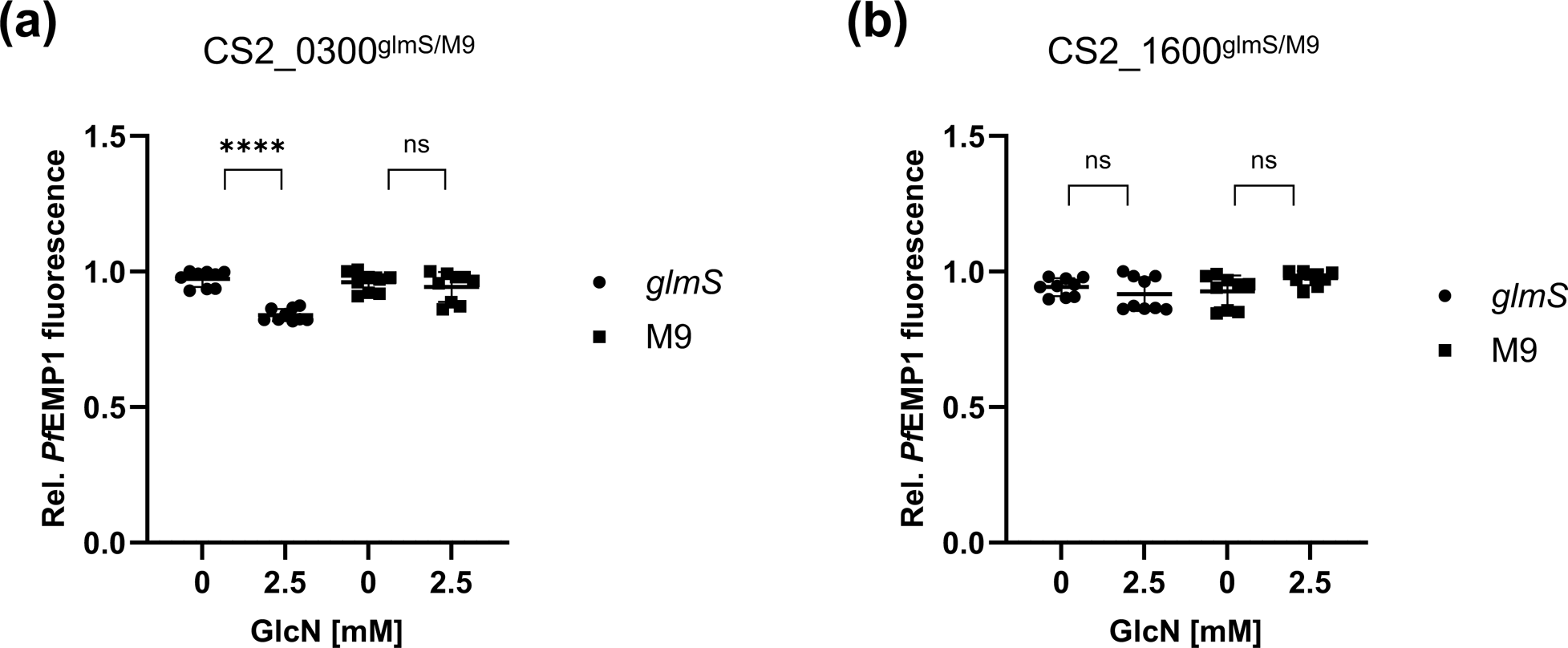
PfEMP1^var2csa^ surface exposure of parasite-infected erythrocytes. Erythrocytes infected with glmS and M9 cell lines, +/− 2.5 mM GlcN treatment for 72 h, were incubated with primary α-PfEMP1^var2csa^ antibody and Cy3-conjugated secondary antibody and PfEMP1^var2csa^ surface exposure was analysed by flow cytometry. Relative PfEMP1 fluorescence signals were statistically evaluated with GraphPad Prism. N=3; ns, not significant; **** p<0.0001 (Students t-test).

## 3. DISCUSSION

Despite initial scepsis, it is now accepted that malaria parasites export a large number of proteins to their host cell, and that these proteins are involved in a number of processes that may be essential for parasite survival either *in vitro* or indeed *in vivo*. A landmark in our understanding of the function of these proteins was placed by Maier et al. in 2008 when they revealed that knockouts of a number of the genes encoding exported proteins led to various aberrations in the ability of the parasite to modify the host cell (Maier et al., 2008). Most importantly, and supported by subsequent studies, many exported proteins appear to be involved in processes underpinning the parasite’s ability to export variant antigens (especially PfEMP1) to the cell surface and present such proteins in a way that allows them to play a role in cytoadherence and likely antigenic variation (Maier et al., 2008; McHugh et al., 2020; Charnaud et al., 2017; Diehl et al., 2021). However, the function of a further number of genes encoding essential exported proteins has been hampered by the lack of the necessary tools for genetic manipulation, making it impossible to analyse genes which fulfil a critical function in blood stage parasites. Over the past few years, a number of systems have been introduced to speed up generation of integrant/knockout parasite lines and allow inducible reductions in protein abundance either by reducing gene expression or by altering the stability of specific mRNAs or the final protein itself. In this current study we utilised the SLI/SLI-TGD approach introduced by Birnbaum and colleagues (Birnbaum et al., 2017), paired with the glmS ribozyme system of Promanna and coworkers (Prommana et al., 2013) to target a number of genes predicted to be exported and involved in host cell modification. Given that a previous study using traditional techniques was able to knockout 77% of the genes of interest (GOI) encoding exported proteins (Maier et al., 2008), and a further study reported a knockout of 61% of all genes targeted using SLI-TGD (Birnbaum et al., 2017), we were quite surprised that our initial SLI-TGD screen did not allow generation of any knockout cell lines, especially as this technique has previously been shown in our laboratory to be able to knockout a gene previously thought to be essential (Maier et al., 2008; Diehl et al., 2021). As a control for both reagents and experimental conditions, we repeated our previous knockout using the vector backbone utilised in this current study and verified that our failure to produce knockouts of the GOIs was not due to technical issues. This further supports an important biological function of the studied GOIs for blood stage growth, and thus reflecting a negatively serendipitous choice of target genes. Introducing a c-terminal GFP-tag via SLI has also been demonstrated to have a high success rate (Birnbaum et al., 2017), however in our hands this technique was only able to generate 2 transgenic parasite lines. Usage of a smaller c-terminal 3xHA-tag did allow isolation of a number of further parasite lines including ones that did not tolerate GFP at this position, suggesting that the large (27 kDa) GFP-tag may have a negative effect on the function of a significant number of the POIs. Taken together, our data are not in direct concordance with previous studies on the successes of various genetic techniques, however we wish to note that the low sample size of all previous studies, and this current one, do not allow any statistical conclusions to be drawn.

For 3 of the 4 POI investigated, we were able to verify export to the host cell. PF^0300^ and PF^1600^ localised to fluorescent foci distributed within the host cell but which later were found not to be Maurer’s clefts (as defined by co-localisation with the MC marker PfSBP1). Such fluorescent structures may represent J-dots, K-dots or Maurer’s cleft tethers (Pachlatko et al., 2010; Kats et al., 2014; Külzer et al., 2010), however a more detailed analysis will be required to corroborate this. While PF^0300^ is predicted to contain a hydrophobic transmembrane domain which may lead to membrane integration, PF^1600^ has no obvious membrane binding motif such as those identified in PfMAHRP or PfKAHRP, but may be recruited to membranes by association with a further (as yet unknown) membrane protein (Rug et al., 2006; Wickham et al., 2001; Spycher et al., 2008). PF^0600^ seemed to localise to the inner leaflet of the erythrocyte plasma membrane in a similar pattern to that observed for the knob protein PfKAHRP (Rug et al., 2006; Wickham et al., 2001) and therefore PF^0600^ may represent a novel knob constituent.

Several publications utilising the glmS ribozyme system in *P. falciparum* demonstrate that the efficiency of knockdown is highly gene dependent, potentially due to different levels of gene expression and thus mRNA concentration (Boddey et al., 2010; Prommana et al., 2013; Elsworth et al., 2014). In our hands, all 4 cell lines showed a significant reduction in protein abundance upon addition of GlcN, although the efficiency differed slightly between cell lines. None of the glmS knockdown parasite lines showed a significant growth disadvantage upon addition of GlcN. This is surprising as our initial SLI-TGD experiments failed to generate any knockout parasite lines, and we were also not able to introduce the GFP coding sequence into these genes, both lines of evidence thus suggesting essentiality of the GOI. Potentially the level of knockdown in all lines was not sufficient to reveal any growth defects or this observation may be due to limitations in the technique itself. We do however wish to note again that control experiments largely verified the genetic approach we used. Consistent with these data, we could observe no reduction in merozoite numbers or a shift in the timing of the developmental cycle.

Previous publications have shown that, even despite normal parasite growth, knockout or knockdown of a number of genes encoding exported proteins can nevertheless lead to aberrations in the parasite’s ability to alter its host cell (Maier et al., 2008; McHugh et al., 2020; Charnaud et al., 2017; Diehl et al., 2021). In this current study, we could not observe any obvious changes either in the distribution of a number of further exported proteins, nor any apparent mutant phenotypes by TEM. In 3 of the 4 cell lines investigated, we did discern a highly statistically significant drop in parasite ability to adhere to CSA in a static binding assay. This cannot be due to a slowing down of the parasites’ developmental cycle upon GlcN treatment (Charnaud et al., 2017), as Giemsa-stained slides showed no difference between treated and untreated lines. Therefore, this drop in cytoadherence suggests a block in the parasite’s ability to either traffic or present functional PfEMP1^var2csa^ to/at the surface of the host cell. To distinguish between a block in transport and attenuation of correct folding or presentation at the cell surface, we detected surface exposed PfEMP1^var2csa^ using specific antisera. This revealed that a knockdown of PF^0300^ led to significantly reduced surface exposure of PfEMP1^var2csa^ compared to controls. In contrast, knockdown of PF^1600^ does not affect surface exposure, despite this cell line showing a more significant drop in CSA binding upon GlcN treatment. As we have previously demonstrated, drops in cytoadherence do not necessarily directly correlate with drops in surface exposure of PfEMP1 (Diehl et al., 2021). In that case, we suspected that lack of CSA binding ability was a product of both, reduced PfEMP1 exposure and abnormally structured knobs. In this current case, however, we could not detect deformation in knob formation, suggesting that this physico-mechanical explanation is unlikely. This must lead us to conclude that either surface-exposed PfEMP1 is not correctly folded and thus cannot bind CSA, or that even a minimal drop in surface exposure of PfEMP1 can dramatically affect static binding ability. At this time, we cannot distinguish between these two explanations.

Our study reveals 3 further exported proteins involved in the cytoadherence phenomenon. We believe that our data, taken together with those from multiple other studies from others and ourselves, also begins to finally shed light on the complexity of the extracellular protein transport system installed by the parasite into the host cell. Despite the identification of numerous proteins required for correct transport and presentation of PfEMP1 on the cell surface, few attempts have been made to understand the precise molecular mechanisms involved in this process. Specifically, we still do not know which exported proteins are involved directly (i.e. by binding to PfEMP1 and directing its traffic), or indirectly (i.e. by being involved in the generation and maintenance of extra-parasitic structures such as knobs and Maurer’s clefts required for traffic or presentation of PfEMP1). Examples from other systems suggest that such trafficking pathways are mediated not just by individual proteins, but by dynamic complexes of multiple molecular players (Barlowe and Miller, 2013). Only once we begin to understand the precise molecular interplay between the numerous exported proteins, will we finally gain insight into this novel process and identify key molecules involved, which may form the basis for potential interventions.

## 4. EXPERIMENTAL PROCEDURES

### 4.1 Generation of SLI-TGD constructs

Fragments encoding homology regions required for integration were amplified using the primers listed in Supplementary 3 and cloned into the pSLI-TGD vector (Birnbaum et al., 2017) or pSLI-glmS/M9-GFP or -HA (a gift of Markus Ganter) using the NotI and MluI sites. All final constructs were verified by sequencing.

### 4.2 Transfection of CS2 parasites

The parental CS2 line had been selected on CSA to ensure expression of the *var2csa* gene and thus correct binding phenotype. Transfections were carried out by electroporation using 150 µg of plasmid DNA according to standard protocols (Fidock and Wellems, 1997) followed by initial selection using 2.5 nM WR99210. For SLI, parasites were then selected with 400-800 µg/ml G418 (Birnbaum et al., 2017). If no parasites were seen following second selection after 10 weeks (3 biological replicates), the integration was deemed to have failed. Following renewed parasite growth, single clones were obtained by limiting dilution and used for further analysis.

### 4.3 Verification of correct integration

Correct integration at the 5’ and 3’ end was verified by PCR on genomic DNA using the primers listed in Supplementary 3. Multiple integration bands were counted as incorrect integration and parasites were excluded from further analysis.

### 4.4 Verification of intact chromosome 2

Parasites were regularly selected using gelatine to ensure a “knobby” phenotype. To detect any deletion of the sub-telomeric region of chromosome 2 which may lead to experimental artefacts, we amplified segments from the *kahsp40* (PlasmoDB: PF3D7_0201800), *pfemp3* (PF3D7_0201900) and *kahrp* (PF3D7_0202000) genes using the primers listed in Supplementary 3. To further check the identity of PCR products, they were treated with HindIII (*kahsp40*), AluI (*pfemp3*) or NdeI (*kahrp*). All parasite clones showed intact telomeric ends.

### 4.5 Immunofluorescence and fluorescence microscopy

Fixation was carried out either according to Tonkin *et al*. (Tonkin et al., 2004) or on glass slides after fixation with acetone/10% methanol. Samples were blocked in 3% BSA/PBS for 2 h and primary antibody (in 3% BSA/PBS) added overnight. Samples were then washed 3×10 min in PBS and secondary antibody (in 3% BSA/PBS) added for 2 h at RT. Samples were then rewashed 3×10 min in PBS and stained with DAPI (1 µg/ml in PBS) for visualisation of the nuclear DNA. Imaging was carried out on a Zeiss Cell Observer epifluorescence microscope equipped with the necessary filter sets, images were captured using Zen (blue) and processed using FiJi. All images were processed and adjusted according to the recommendations of (Rossner and Yamada, 2004). All primary and secondary antibodies and concentrations are listed in Supplementary 3.

### 4.6 Western blot and immunodetection

Samples representing 1×10^7^ infected erythrocytes (isolated by MACS) were prepared by boiling in Laemmli sample buffer and separated on 10-15% SDS-PAGE. Transfer to nitrocellulose membrane was carried out at 25 V and 1 A for 30 min in a Trans-Blot Turbo system (Bio-Rad, Hercules, USA). Membranes were blocked in 5% milk powder/PBS for 1 h, followed by incubation with primary antibodies diluted in blocking solution overnight. Membranes were washed 3×10 min in PBS and secondary antibodies diluted in blocking solution added for 2 h. Following 3×10 min wash in PBS, bound antibodies were visualised using ECL/H_2_O_2_ in an Intas ECL ChemoStar. Exposure times were selected to avoid image saturation. Band quantification was carried out using FiJi and normalised against the aldolase loading control. All primary and secondary antibodies and concentrations are listed in Supplementary 3.

### 4.7 Transmission electron microscopy

Trophozoite-stage iRBC were enriched by gelatine flotation and fixed in 100 mM cacodylate solution (CaCo) with 2% glutaraldehyde and 2% paraformaldehyde overnight at 4 °C. Cells were then washed 3×5 min with CaCo solution and incubated with 1% osmium in CaCo solution for 1 h at RT. After 2x washing with ddH2O, the cells were incubated in 1% U-acetate in ddH2O overnight at 4 °C to increase the contrast in the final image. The next day, the cells were washed 2×10 min ddH2O at RT and dehydrated in a series of 25%, 50%, 70%, 90%, and 2×100% acetone for 10 min each. Following this, cells were incubated in a series of 25%, 50%, and 75% Spurr for 45 min each at RT and stored in 100% Spurr overnight before the sections were prepared and imaged on a Jeol JEM-1400 electron microscope at 80 kV.

### 4.8 CSA Binding assay

The CSA binding assay was carried out according to standard protocols (Beeson et al., 1999), including a soluble CSA specificity control. Images were captured using a Kern OCM inverted microscope and binding determined using a previously developed algorithm in Ilastik and ImageJ (Diehl et al., 2021).

### 4.9 Detection of PfEMP1^var2csa^ on the erythrocyte surface

Infected erythrocytes were isolated using MACS and incubated with rabbit anti-var2CSA antisera (11P, a kind gift of Benoit Gamain) in 0.3% BSA/PBS for 30 min at 4 °C. Samples were washed 3×10 min in PBS followed by incubation with anti-rabbit Cy3 secondary antibody for 30 min at 4 °C. Following further washing, samples were fixed in 4% paraformaldehyde/PBS overnight. Subsequently, samples were washed 3×10 min in PBS, stained with 1 µg/ml DAPI/PBS for 20 min and washed 3×10 min in PBS. Samples were measured in a CytoflexS using the corresponding laser lines, and analysis carried out using the Cytoflex software.

### 4.10 Parasite growth and development assay

Parasite growth was determined using SybrGreen (Bennett et al., 2004; Smilkstein et al., 2004), an initial parasitaemia (ring stages) of 0.2%, and assay length of 72 h. In parallel, slides were prepared at the time points noted, stained with Giemsa and DIC images captured. Merozoites were counted from Giemsa-stained images at 78 hpi.

### 4.11 Treatment of glmS/M9 lines with GlcN

Parasites were synchronised to different stages depending on the experiment (see above for details), split into parallel cultures and GlcN (Sigma) added for 72 h.

### 4.12 Parasite culture

Parasites were cultured in A^+^ erythrocytes (haematocrit 4%, blood bank Gießen) and complete RPMI/10% human serum (blood bank Gießen) in a gassed incubator at standard conditions. Parasite growth was monitored by Giemsa-stained slides and maintained at <10% parasitaemia.

## ACKNOWLEDGMENTS

This project was financed by the DFG by a grant to JMP (Project number 351262938). We thank numerous colleagues for providing materials and protocols.

## AUTHOR CONTRIBUTIONS

**Nina Küster:** Investigation; methodology; writing – review and editing; visualization. **Lena Roling**: Investigation; methodology. **Ardin Ouayoue**: Investigation; methodology. **Katharina Steeg**: Investigation. **Jude M Przyborski**: Writing – original draft; writing – review and editing; investigation; supervision; funding acquisition; conceptualization; project administration.

## ABBREVIATED SUMMARY

We have identified a number of proteins involved in host cell modification by malaria parasites. Furthermore, we highlight a number of genes which appear to be essential for blood stage survival of these parasites.

## Supplementary Figure Legends

**S1. Sequence analysis.**

**S2. Generation of glmS knockdown cell lines.** (Top) Strategy to generate glmS knockdown cell lines via SLI. The native genomic locus was modified to add an additional c-terminal skip-peptide (S) and HA-tag or GFP-tag to the GOI using homologous recombination and selection-linked integration (SLI). UTR, untranslated region. Primers used for integration PCRs are indicated by arrows. (Lower panels) Verification of M9- and glmS knockdown cell lines via Integration-PCR. GlmS and M9 cell lines were tested for 5’ (primer A and B) and 3’ integration (primer C and D), episomal plasmids (primer C and B), and wildtype-locus (primer A and D). CS2, WT-positive control; ddH20, negative control. Additionally, integrity of the subtelomeric regions of chromosome 2 which harbours *kahsp40, emp3* and *kahrp* was verified by PCR and restriction digest.

**S3. List of primers and antibodies.**

**S4. Middle section of deconvoluted Z-stack from Figure 1.**

**S5. Deconvoluted Z-stack Movie from Figure 1.**

**S6. Light microscopy of parasite-infected erythrocytes.** Cell morphology and cell cycle progression of glmS and M9 cell lines was monitored for 72 h by imaging Giemsa-stained blood smears of parasites at the time points and with the GlcN concentrations indicated. Scale bar, 5 µm. All images are representative of at least 10 independent observations.

**S7. Localisation of other exported proteins in M9 cell lines.** Immunofluorescent localisation of PfKAHRP, PfSBP1, PfREX3 and PfEMP3 in M9 cell lines +/− 2.5 mM GlcN treatment for 72 h. Cells were fixed and incubated with anti-PfKAHRP, anti-PfSBP1, anti-PfREX3 or anti-PfEMP3 respectively followed by Cy3-conjugated secondary antibody. Fluorescence channels are shown individually in black/white for highest contrast. DIC, differential interference contrast; DAPI, nucleus. All images are representative of at least 20 independent observations.

## LEGEND GRAPHICAL ABSTRACT

***In silico* workflow for selection of genes targeted in this study.**

