## Supplementary 1 for "A systematic targeted genetic screen identifies proteins involved in cytoadherence of the malaria parasite *P. falciparum*"

**Predicted protein sequences POI**
Predicted TM domain in blue
PEXEL sequence in red

>300
MNMFFLFIKIFIFSIFTINLKLTNRNDYNIPYKGKKNLLGKRLGLISCRTLTEVYDAFNDATVKMLDISFDSSKASTTKT 25kDa pI 10.58
RRFIHDRYSTAPYELIKRKPKKKKSPFSKMKVKFVRYFQKINYILNEILYFFMDNTNCFFLPTKYILAPFVFTFKILYDI
IAFIAIVFILVIIIIYTLIKKMCLRMHYSDSMQNLRSRKNHEAKVKIYKPEA

>600
MGYSNNKFNIFTLWNNIILYFILIVTFTFYNKDLFNKYNGEKSNIGASFNFGNNRSLAEYYNNKDGYNVLRVNLDHKNLK 26kDa pI 9.83
DVLGNMHPEIKMVEVDSESVCPGTNEVNLKVVTNIPPDMIKVNATSENMSVGQWDYIMQYYGQSTPKEVSKLDSEVKDKI
EKKIKKKKRKTPLIRYIAELVGYGIIFIPGFPVLVGIVSVGFCILIFMGKKSAKNYFSTIKKWLF

>1600
MITKNYNISKNVIRQIGIGVNKSICFTCKNINILGKCQKICIKHFLIFNKILLLFILVWIFQYSNEEFTCKGSRSTKQDE 88kDa pI 4.77
EKASLRFYRLLAQSYVFNSRLDGFHNYSNAENERMFMRNYMNDGLFKSQAYNESEKRDNLRNNIYNEQNDFAKEGYSREY
DSKYNSAEFGTSRKKNDGRNPFNYQSNNENNKPASYMNEDIYNKNYYNQRENIDELNKKLHDDWSNFLNEDYSNEFESNN
FNSPHYDTYKSEYDTYKSEYDTYKSQYDTYKSQYDDRNKYSYPSREGNYVPPFMDENDNRLNNNYRSGFDDTKKNNINDD
KGMNFVRDEFYDEYNNNNMNTSPEYMFSQEWDNQNTHDGFPSGWNNYNSENSDHKDQIKVETPYIRVVEEVNDESNNKRD
NYDTLNKDSSCNRNEKMEYYFQSDKFPVDCKGIYTNSNIIVEEYDTPVDLKETNDNFYSFNEKLNNERQKKYMREDKEFI
NEMGHMSSTSSNSPSSHFPSPHSQPSFIPETFQDSNIESPEFELPTFESATVHTSEYELPNFDILEYDLPHFDPSEFKPE
IFDESEFKPEIFDESEFKPSVFDPSEYKNSPFKESKISTSSHSTRSYENEFNLSTDRRNSESSSRIHVDDENYKRLKKIL
SNLCSSREKIYDLSKEEKEFLVKMLQLDYEDFNFNYRGNTRSYNDSSPFTNFELKRIHLKIKAFIYKCLLRLLQSYEKST
ESRVGSNIYDIY

>3300
MKNIKNMKNIKSEGFIFFVLVFFICIIFGCIYESLHEGPYKKTLNSLHESTKYRHFNKIRLLTEYKDTLQIKVEQKSLRD 71kDa pI 7.98
YVNNDRYNNVNTNDYTSYKDKGEQFNDTICVVDKKKENVTINNEEECNKNFYQYLQYLEHNNKQDNKYEETNYFLQGNDK
HIDSEHNGINKMYKETIHKTLTSDVSTENSYTHNNSRDDEPQNGKRTYNNQSNNNLPYDNSSYNISPYHGPNNNVPYNKS
NNFEQCNTQDNKHCNDLDTYHTCYGPDNYPQKYDNYRQECDNYRQECDNYRQEYDNYPQKYDNYRQECDNYRQEYDNYPH
GFDNYPRGFDNYPHGYDNHPHRPHIYPHGFDNHPHRPHMYPHNFPMRNESVGGPYYRPPHIIERSNYYKNPKKAPHNMML
PCDTMKDNKSICDEQNFQRELEKIIKKNNLQNGNIRDNHDTRINDYNKRLTEYNKRLTEYNKRLTEYTKRLNEHYKRNGY
NIQNRQNSIERAQSNDVVLYGHNFQNAFRYKQNTRSYYPHVNSNEATHHQKTMYFTQQNNYSREEYPIKSEQHLYHVKSK
RLEKKLYDYQNGTNPVTNFLERHF

**Alignment Syntenic Orthologues PF3D7_0301600**

```
PADL01_0034200-t36_1-p1         MINKNYNISKNGIRQIGIRANMTICFTSKHINIIGKCQKKYIKHFLIFNKVLLLFVLVCI    60
PGABG01_0300800-t36_1-p1        MINKNYNISKNGIRQIGIRANRTICFTSKHINIIGKCQKKYIKHFLIFNKFLLLFILVSI    60
PGSY75_0301600-t31_1-p1         MINKNYNISKNGIRQIGIRANRTICFTSKHINIIGKCQKKYIKHFLIFNKVLLLFILVCI    60
PBLACG01_0301900-t36_1-p1       MITQNYNISKNVIRQIGIRANRSICFTCKHINILQKCQKIYIKHFLIFNKILLLFILVWI    60
PRG01_0304500-t36_1-p1          MVTKNYNISKNVIRQIGIGVNKSICFTFKNINILGKCQKICIKHFLIFNKILLLFILVWI    60
PRCDC_0300700.1-p1              MVTKNYNISKNVIRQIGIGVNKSICFTFKNINILGKCQKICIKHFLIFNKILLLFILVWI    60
PfML01_030006900-t41_1-p1       MITKNYNISKNVIRQIGIGVNKSICFTCKNINILGKCQKICIKHFLIFNKILLLFILVWI    60
PfGB4_030007500-t41_1-p1        MITKNYNISKNVIRQIGIGVNKSICFTCKNINILGKCQKICIKHFLIFNKILLLFILVWI    60
PfSD01_030006800-t41_1-p1       MITKNYNISKNVIRQIGIGVNKSICFTCKNINILGKCQKICIKHFLIFNKILLLFILVWI    60
PfKH02_030007200-t41_1-p1       MITKNYNISKNVIRQIGIGVNKSICFTCKNINILGKCQKICIKHFLIFNKILLLFILVWI    60
PfIT_030006400-t41_1-p1         MITKNYNISKNVIRQIGIGVNKSICFTCKNINILGKCQKICIKHFLIFNKILLLFILVWI    60
PfKH01_030006400-t41_1-p1       MITKNYNISKNVIRQIGIGVNKSICFTCKNINILGKCQKICIKHFLIFNKILLLFILVWI    60
PfNF135_030006800.1-p1          MITKNYNISKNVIRQIGIGVNKSICFTCKNINILGKCQKICIKHFLIFNKILLLFILVWI    60
PfCD01_030007000-t41_1-p1       MITKNYNISKNVIRQIGIGVNKSICFTCKNINILGKCQKICIKHFLIFNKILLLFILVWI    60
PfGN01_030007300-t41_1-p1       MITKNYNISKNVIRQIGIGVNKSICFTCKNINILGKCQKICIKHFLIFNKILLLFILVWI    60
PfKE01_030006300-t41_1-p1       MITKNYNISKNVIRQIGIGVNKSICFTCKNINILGKCQKICIKHFLIFNKILLLFILVWI    60
PfSN01_030007100-t41_1-p1       MITKNYNISKNVIRQIGIGVNKSICFTCKNINILGKCQKICIKHFLIFNKILLLFILVWI    60
PfTG01_030008300-t41_1-p1       MITKNYNISKNVIRQIGIGVNKSICFTCKNINILGKCQKICIKHFLIFNKILLLFILVWI    60
Pf7G8-2_000071200.1-p1          MITKNYNISKNVIRQIGIGVNKSICFTCKNINILGKCQKICIKHFLIFNKILLLFILVWI    60
Pf7G8_030007400-t41_1-p1        MITKNYNISKNVIRQIGIGVNKSICFTCKNINILGKCQKICIKHFLIFNKILLLFILVWI    60
PfHB3_030005500-t41_1-p1        MITKNYNISKNVIRQIGIGVNKSICFTCKNINILGKCQKICIKHFLIFNKILLLFILVWI    60
PF3D7_0301600.1-p1              MITKNYNISKNVIRQIGIGVNKSICFTCKNINILGKCQKICIKHFLIFNKILLLFILVWI    60
PfNF54_030006800.1-p1           MITKNYNISKNVIRQIGIGVNKSICFTCKNINILGKCQKICIKHFLIFNKILLLFILVWI    60
PfGA01_030008000-t41_1-p1       MITKNYNISKNVIRQIGIGVNKSICFTCKNINILGKCQKICIKHFLIFNKILLLFILVWI    60
PfDd2_030006600-t41_1-p1        MITKNYNISKNVIRQIGIGVNKSICFTCKNINILGKCQKICIKHFLIFNKILLLFILVWI    60
                                *:.:******* ****** .* :**** *:***: ****  *********.****:** *

PADL01_0034200-t36_1-p1         CQYSIDEITFNGSRSIKQDEEKASLRFNRLLAQSFVFNSRLDGSYNYSNVENERMYMPHY    120
PGABG01_0300800-t36_1-p1        CQYSIDEITFNGSRSTKQDEEKASLRFNRLLAQSFVFNSRLDGSYNYPNVENERMYMPHY    120
PGSY75_0301600-t31_1-p1         YQYSIDEITFNGSRSTKQDEEKASLRFNRLLAQSFVFNSRLDGSYNYPNVENERMYMPHY    120
PBLACG01_0301900-t36_1-p1       FQYSNEDFNCKGSRSTKHDEERASLRFYRLLAQSYIFNSRLDGSYNYSNVENERMYMRDY    120
PRG01_0304500-t36_1-p1          FQYSNEEFTCKGSRSTKQDEEKASLRFYRLLAQSYLFNSRLDGSYNYSNAENERMFMRNY    120
PRCDC_0300700.1-p1              FQYSNEEFTCKGSRSTKQDEEKASLRFYRLLAQSYLFNSRLDGSYNYSNAENERMFMRNY    120
PfML01_030006900-t41_1-p1       FQYSNEEFTCKGSRSTKQDEEKASLRFYRLLAQSYVFNSRLDGFHNYSNAENERMFMRNY    120
PfGB4_030007500-t41_1-p1        FQYSNEEFTCKGSRSTKQDEEKASLRFYRLLAQSYVFNSRLDGFHNYSNAENERMFMRNY    120
PfSD01_030006800-t41_1-p1       FQYSNEEFTCKGSRSTKQDEEKASLRFYRLLAQSYVFNSRLDGFHNYSNAENERMFMRNY    120
PfKH02_030007200-t41_1-p1       FQYSNEEFTCKGSRSTKQDEEKASLRFYRLLAQSYVFNSRLDGSHNYSNAENERMFMRNY    120
PfIT_030006400-t41_1-p1         FQYSNEEFTCKGSRSTKQDEEKASLRFYRLLAQSYVFNSRLDGSHNYSNAENERMFMRNY    120
PfKH01_030006400-t41_1-p1       FQYSNEEFTCKGSRSTKQDEEKASLRFYRLLAQSYVFNSRLDGSHNYSNAENERMFMRNY    120
PfNF135_030006800.1-p1          FQYSNEEFTCKGSRSTKQDEEKASLRFYRLLAQSYVFNSRLDGFHNYSNAENERMFMRNY    120
PfCD01_030007000-t41_1-p1       FQYSNEEFTCKGSRSTKQDEEKASLRFYRLLAQSYVFNSRLDGFHNYSNAENERMFMRNY    120
PfGN01_030007300-t41_1-p1       FQYSNEEFTCKGSRSTKQDEEKASLRFYRLLAQSYVFNSRLDGFHNYSNAENERMFMRNY    120
PfKE01_030006300-t41_1-p1       FQYSNEEFTCKGSRSTKQDEEKASLRFYRLLAQSYVFNSRLDGFHNYSNAENERMFMRNY    120
PfSN01_030007100-t41_1-p1       FQYSNEEFTCKGSRSTKQDEEKASLRFYRLLAQSYVFNSRLDGFHNYSNAENERMFMRNY    120
PfTG01_030008300-t41_1-p1       FQYSNEEFTCKGSRSTKQDEEKASLRFYRLLAQSYVFNSRLDGFHNYSNAENERMFMRNY    120
Pf7G8-2_000071200.1-p1          FQYSNEEFTCKGSRSTKQDEEKASLRFYRLLAQSYVFNSRLDGFHNYSNAENERMFMRNY    120
Pf7G8_030007400-t41_1-p1        FQYSNEEFTCKGSRSTKQDEEKASLRFYRLLAQSYVFNSRLDGFHNYSNAENERMFMRNY    120
PfHB3_030005500-t41_1-p1        FQYSNEEFTCKGSRSTKQDEEKASLRFYRLLAQSYVFNSRLDGFHNYSNAENERMFMRNY    120
PF3D7_0301600.1-p1              FQYSNEEFTCKGSRSTKQDEEKASLRFYRLLAQSYVFNSRLDGFHNYSNAENERMFMRNY    120
PfNF54_030006800.1-p1           FQYSNEEFTCKGSRSTKQDEEKASLRFYRLLAQSYVFNSRLDGFHNYSNAENERMFMRNY    120
PfGA01_030008000-t41_1-p1       FQYSNEEFTCKGSRSTKQDEEKASLRFYRLLAQSYVFNSRLDGFHNYSNAENERMFMRNY    120
PfDd2_030006600-t41_1-p1        FQYSNEEFTCKGSRSTKQDEEKASLRFYRLLAQSYVFNSRLDGSHNYSNAENERMFMRNY    120
                                 *** :::. :**** *:***:***** ******::******* :** *.*****:* .*
```

```
PADL01_0034200-t36_1-p1        MNNNFFKSQAYNESEKRDNLRSKIYNEQNDFAKEGYSREYDSKYNSQEFGTSRKKNNGRN    180
PGABG01_0300800-t36_1-p1       MNNNFFKSQAYNESEKRDNLRSKIYNEQNDFPKEGYSREYDSKYNSQEFGTSRKKNNGRN    180
PGSY75_0301600-t31_1-p1        MNNNFFKSQAYNESEKRDNLRSKIYNEQNNFPKEGYSREYDSKYNSQEFGTSRKKNNGRN    180
PBLACG01_0301900-t36_1-p1      MNNNLFKSQAYNESEKRDNLRDNIYNEKNDFAKEGYSREYDSKCNSPEFGTSRKKNDGRN    180
PRG01_0304500-t36_1-p1         MNDGLFKSQAYNESEKRDNLRNNIYNEQNDFAKEGYSREYDSKFNSAEFGTSRKKNGGRN    180
PRCDC_0300700.1-p1             MNDGLFKSQAYNESEKRDNLRNNIYNEQNDFAKEGYSREYDSKFNSAEFGTSRKKNGGRN    180
PfML01_030006900-t41_1-p1      MNDGLFKSQAYNESEKRDNLRNNIYNEQNDFAKEGYSREYDSKYNSAEFGTSRKKNDGRN    180
PfGB4_030007500-t41_1-p1       MNDGLFKSQAYNESEKRDNLRNNIYNEQNDFAKEGYSREYDSKYNSAEFGTSRKKNDGRN    180
PfSD01_030006800-t41_1-p1      MNDGLFKSQAYNESEKRDNLRNNIYNEQNDFAKEGYSREYDSKYNSAEFGTSRKKNDGRN    180
PfKH02_030007200-t41_1-p1      MNDGLFKSQAYNESEKRDNLRNNIYNEQNDFAKEGYSREYDSKYNSAEFGTSRKKNDGKN    180
PfIT_030006400-t41_1-p1        MNDGLFKSQAYNESEKRDNLRNNIYNEQNDFAKEGYSREYDSKYNSAEFGTSRKKNDGKN    180
PfKH01_030006400-t41_1-p1      MNDGLFKSQAYNESEKRDNLRNNIYNEQNDFAKEGYSREYDSKYNSAEFGTSRKKNDGKN    180
PfNF135_030006800.1-p1         MNDGLFKSQAYNESEKRDNLRNNIYNEQNDFAKEGYSREYDSKYNSAEFGTSRKKNDGKN    180
PfCD01_030007000-t41_1-p1      MNDGLFKSQAYNESEKRDNLRNNIYNEQNDFAKEGYSREYDSKYNSAEFGTSRKKNDGRN    180
PfGN01_030007300-t41_1-p1      MNDGLFKSQAYNESEKRDNLRNNIYNEQNDFAKEGYSREYDSKYNSAEFGTSRKKNDGRN    180
PfKE01_030006300-t41_1-p1      MNDGLFKSQAYNESEKRDNLRNNIYNEQNDFAKEGYSREYDSKYNSAEFGTSRKKNDGRN    180
PfSN01_030007100-t41_1-p1      MNDGLFKSQAYNESEKRDNLRNNIYNEQNDFAKEGYSREYDSKYNSAEFGTSRKKNDGRN    180
PfTG01_030008300-t41_1-p1      MNDGLFKSQAYNESEKRDNLRNNIYNEQNDFAKEGYSREYDSKYNSAEFGTSRKKNDGRN    180
Pf7G8-2_000071200.1-p1         MNDGLFKSQAYNESEKRDNLRNNIYNEQNDFAKEGYSREYDSKYNSAEFGTSRKKNDGRN    180
Pf7G8_030007400-t41_1-p1       MNDGLFKSQAYNESEKRDNLRNNIYNEQNDFAKEGYSREYDSKYNSAEFGTSRKKNDGRN    180
PfHB3_030005500-t41_1-p1       MNDGLFKSQAYNESEKRDNLRNNIYNEQNDFAKEGYSREYDSKYNSAEFGTSRKKNDGRN    180
PF3D7_0301600.1-p1             MNDGLFKSQAYNESEKRDNLRNNIYNEQNDFAKEGYSREYDSKYNSAEFGTSRKKNDGRN    180
PfNF54_030006800.1-p1          MNDGLFKSQAYNESEKRDNLRNNIYNEQNDFAKEGYSREYDSKYNSAEFGTSRKKNDGRN    180
PfGA01_030008000-t41_1-p1      MNDGLFKSQAYNESEKRDNLRNNIYNEQNDFAKEGYSREYDSKYNSAEFGTSRKKNDGRN    180
PfDd2_030006600-t41_1-p1       MNDGLFKSQAYNESEKRDNLRNNIYNEQNDFAKEGYSREYDSKYNSAEFGTSRKKNDGKN    180
                               **:.:****************.:****:*:* *********** ** *********.*:*

PADL01_0034200-t36_1-p1        SFNYQSNNENNKHSSYMNEDMYNKNYFNQRENIDELNKKMHDDWSNYLNEDYSHGFESNN    240
PGABG01_0300800-t36_1-p1       SFNYQSNNENNKHSFYMNEDMYNKNYFDQRENIDELNKKMHDDWSNFLIEDYSHEFESNN    240
PGSY75_0301600-t31_1-p1        SFNYQSNNENNKHSFYMNEDMYNKNYFDQRENIDELNKKMHDDWSNFLIEDYSHEFESNN    240
PBLACG01_0301900-t36_1-p1      SFNYQSNNENNKPSSYMNEDIYNKNYYEQRENIDEINKKLHDDWSNFLNENYSSEFESNN    240
PRG01_0304500-t36_1-p1         PFNYQSNNENNKPASFMNEDIYNKNYYNERENIDELNKKLHDDWSKFLNEDYSNEFESNN    240
PRCDC_0300700.1-p1             PFNYQSNNENNKPASFMNEDIYNKNYYNERENIDELNKKLHDDWSKFLNEDYSNEFESNN    240
PfML01_030006900-t41_1-p1      PFNYQSNNENNKPASYMNEDIYNKNYYNQRENIDELNKKLHDDWSNFLNEDYSNEFESNN    240
PfGB4_030007500-t41_1-p1       PFNYQSNNENNKPASYMNEDIYNKNYYNQRENIDELNKKLHDDWSNFLNEDYSNEFESNN    240
PfSD01_030006800-t41_1-p1      PFNYQSNNENNKPASYMNEDIYNKNYYNQRENIDELNKKLHDDWSNFLNEDYSNEFESNN    240
PfKH02_030007200-t41_1-p1      PFNYQSNNENNKPASYMNEDIYNKNYYNQRENIDELNKKLHDDWSNFLNEDYSNEFESNN    240
PfIT_030006400-t41_1-p1        PFNYQSNNENNKPASYMNEDIYNKNYYNQRENIDELNKKLHDDWSNFLNEDYSNEFESNN    240
PfKH01_030006400-t41_1-p1      PFNYQSNNENNKPASYMNEDIYNKNYYNQRENIDELNKKLHDDWSNFLNEDYSNEFESNN    240
PfNF135_030006800.1-p1         PFNYQSNNENNKPASYMNEDIYNKNYYNQRENIDELNKKLHDDWSNFLNEDYSNEFESNN    240
PfCD01_030007000-t41_1-p1      PFNYQSNNENNKPASYMNEDIYNKNYYNQRENIDELNKKLHDDWSNFLNEDYSNEFESNN    240
PfGN01_030007300-t41_1-p1      PFNYQSNNENNKPASYMNEDIYNKNYYNQRENIDELNKKLHDDWSNFLNEDYSNEFESNN    240
PfKE01_030006300-t41_1-p1      PFNYQSNNENNKPASYMNEDIYNKNYYNQRENIDELNKKLHDDWSNFLNEDYSNEFESNN    240
PfSN01_030007100-t41_1-p1      PFNYQSNNENNKPASYMNEDIYNKNYYNQRENIDELNKKLHDDWSNFLNEDYSNEFESNN    240
PfTG01_030008300-t41_1-p1      PFNYQSNNENNKPASYMNEDIYNKNYYNQRENIDELNKKLHDDWSNFLNEDYSNEFESNN    240
Pf7G8-2_000071200.1-p1         PFNYQSNNENNKPASYMNEDIYNKNYYNQRENIDELNKKLHDDWSNFLNEDYSNEFESNN    240
Pf7G8_030007400-t41_1-p1       PFNYQSNNENNKPASYMNEDIYNKNYYNQRENIDELNKKLHDDWSNFLNEDYSNEFESNN    240
PfHB3_030005500-t41_1-p1       PFNYQSNNENNKPASYMNEDIYNKNYYNQRENIDELNKKLHDDWSNFLNEDYSNEFESNN    240
PF3D7_0301600.1-p1             PFNYQSNNENNKPASYMNEDIYNKNYYNQRENIDELNKKLHDDWSNFLNEDYSNEFESNN    240
PfNF54_030006800.1-p1          PFNYQSNNENNKPASYMNEDIYNKNYYNQRENIDELNKKLHDDWSNFLNEDYSNEFESNN    240
PfGA01_030008000-t41_1-p1      PFNYQSNNENNKPASYMNEDIYNKNYYNQRENIDELNKKLHDDWSNFLNEDYSNEFESNN    240
PfDd2_030006600-t41_1-p1       PFNYQSNNENNKPASYMNEDIYNKNYYNQRENIDELNKKLHDDWSNFLNEDYSNEFESNN    240
                                *********** : :****:*****:::******:***:*****::* *:**  *****
```

```
PADL01_0034200-t36_1-p1        FNSPHYDTYKSQYDTYKSQYDTYKSQYGIYNSQYDTYNSQYDDRNNYSYPSSEENYVPPF    300
PGABG01_0300800-t36_1-p1       FNSPHYDTYKSQYDTYKSQYDTYKSQYGIYNSEYDTYNSEYDDRNNYFYPSSEENFVPPF    300
PGSY75_0301600-t31_1-p1        FNSPHYDTYKSQYDTYKSQYDTYKSQYGIYNSEYDTYNSEYDDRNNYFYPSSEENFVPPF    300
PBLACG01_0301900-t36_1-p1      FNSPHYDTYKSQYDTYKSQYDTYKSQYDNYKSQYETYKSQYDDRNKYSYPSRDENYVPPF    300
PRG01_0304500-t36_1-p1         FNSPHYDTYKSQYDTYKSKYDTYKSPYDTYKSQYDIYKSQYDDRNKYSYPSREANYVPPF    300
PRCDC_0300700.1-p1             FNSPHYDTYKSQYDTYKSKYDTYKSPYDTYKSQYDIYKSQYDDRNKYSYPSREANYVPPF    300
PfML01_030006900-t41_1-p1      FNSPHYDTYKSEYDTYKSEYDTYKSQYDTYKSQYDTYKSQYDDRNKYSYPSREGNYVPPF    300
PfGB4_030007500-t41_1-p1       FNSPHYDTYKSEYDTYKSEYDTYKSQYDTYKSQYDTYKSQYDDRNKYSYPSREGNYVPPF    300
PfSD01_030006800-t41_1-p1      FNSPHYDTYKSEYDTYKSEYDTYKSQYDTYKSQYDTYKSQYDDRNKYSYPSREGNYVPPF    300
PfKH02_030007200-t41_1-p1      FNSPHYDTYKSEYDTYKSEYDT-------YKSQYDTYKSQYDDRNKYSYPSREGNYVPPF    293
PfIT_030006400-t41_1-p1        FNSPHYDTYKSEYDTYKSEYDT-------YKSQYDTYKSQYDDRNKYSYPSREGNYVPPF    293
PfKH01_030006400-t41_1-p1      FNSPHYDTYKSEYDTYKSEYDT-------YKSQYDTYKSQYDDRNKYSYPSREGNYVPPF    293
PfNF135_030006800.1-p1         FNSPHYDTYKSEYDTYKSEYDT-------YKSQYDTYKSQYDDRNKYSYPSREGNYVPPF    293
PfCD01_030007000-t41_1-p1      FNSPHYDTYKSEYDTYKSEYDT-------YKSQYDTYKSQYDDRNKYSYPSREGNYVPPF    293
PfGN01_030007300-t41_1-p1      FNSPHYDTYKSEYDTYKSEYDT-------YKSQYDTYKSQYDDRNKYSYPSREGNYVPPF    293
PfKE01_030006300-t41_1-p1      FNSPHYDTYKSEYDTYKSEYDT-------YKSQYDTYKSQYDDRNKYSYPSREGNYVPPF    293
PfSN01_030007100-t41_1-p1      FNSPHYDTYKSEYDTYKSEYDT-------YKSQYDTYKSQYDDRNKYSYPSREGNYVPPF    293
PfTG01_030008300-t41_1-p1      FNSPHYDTYKSEYDTYKSEYDT-------YKSQYDTYKSQYDDRNKYSYPSREGNYVPPF    293
Pf7G8-2_000071200.1-p1         FNSPHYDTYKSEYDTYKSEYDT-------YKSQYDTYKSQYDDRNKYSYPSREGNYVPPF    293
Pf7G8_030007400-t41_1-p1       FNSPHYDTYKSEYDTYKSEYDT-------YKSQYDTYKSQYDDRNKYSYPSREGNYVPPF    293
PfHB3_030005500-t41_1-p1       FNSPHYDTYKSEYDTYKSEYDT-------YKSQYDTYKSQYDDRNKYSYPSREGNYVPPF    293
PF3D7_0301600.1-p1             FNSPHYDTYKSEYDTYKSEYDT-------YKSQYDTYKSQYDDRNKYSYPSREGNYVPPF    293
PfNF54_030006800.1-p1          FNSPHYDTYKSEYDTYKSEYDT-------YKSQYDTYKSQYDDRNKYSYPSREGNYVPPF    293
PfGA01_030008000-t41_1-p1      FNSPHYDTYKSEYDTYKSEYDT-------YKSQYDTYKSQYDDRNKYSYPSREGNYVPPF    293
PfDd2_030006600-t41_1-p1       FNSPHYDTYKSEYDTYKSEYDT-------YKSQYDTYKSQYDDRNKYSYPSREGNYVPPF    293
                               ***********:******:***       *:*:*: *:*:*****:* *** : *:****

PADL01_0034200-t36_1-p1        MDENMNGLNNNYRTGFDESPKNNINDDEGINFVRDEFYDEYDN-NMNTPPQYMFSPEWDN    359
PGABG01_0300800-t36_1-p1       MDENMNGLNNNYRTGFDESQKNNINDDVGINFVRDEFYDEYNN-NMNTPPQYMFSPEWDN    359
PGSY75_0301600-t31_1-p1        MDENMNGLNNNYRTGFDESQKNNINDDEGINFVRDEFYDEYNN-NMNTPPQYMFSPEWDN    359
PBLACG01_0301900-t36_1-p1      MDENMNRLNNNYRSGFDDSQKNNINDDGGMNFVRDEFYDEYNN-NMNTPPQYMFSPEWDN    359
PRG01_0304500-t36_1-p1         MDENINRLNNNYRDGFDDTKKNNINDDKGMNFVRDEFYDEYNN-NMNTSPEYLFSQEWDN    359
PRCDC_0300700.1-p1             MDENINRLNNNYRDGFDDTKKNNINDDKGMNFVRDEFYDEYNN-NMNTSPEYLFSQEWDN    359
PfML01_030006900-t41_1-p1      MDENDNRLNNNYRSGFDDTKKNNINDDKGMNFVRDEFYDEYNNNNMNTSPEYMFSQEWDN    360
PfGB4_030007500-t41_1-p1       MDENVNRLNNNYKSGFDDTKKNNINDDKGMNFVRDEFYDEYNNNNMNTSPEYMFSQEWDN    360
PfSD01_030006800-t41_1-p1      MDENDNRLNNNYRSGFDDTKKNNINDDKGMNFVRDEFYDEYNNNNMNTSPEYMFSQEWDN    360
PfKH02_030007200-t41_1-p1      MDENVNRLNNNYKSGFDDTKKNNINDDKGMNFVRDEFYDEYNNNNMNTSPEYMFSQEWDN    353
PfIT_030006400-t41_1-p1        MDENDNRLNNNYRSGFDDTKKNNINDDKGMNFVRDEFYDEYNNNNMNTSPEYMFSQEWDN    353
PfKH01_030006400-t41_1-p1      MDENDNRLNNNYRSGFDDTKKNNINDDKGMNFVRDEFYDEYNNNNMNTSPEYMFSQEWDN    353
PfNF135_030006800.1-p1         MDENDNRLNNNYRSGFDDTKKNNINDDKGMNFVRDEFYDEYNNNNMNTSPEYMFSQEWDN    353
PfCD01_030007000-t41_1-p1      MDENDNRLNNNYRSGFDDTKKNNINDDKGMNFVRDEFYDEYNNNNMNTSPEYMFSQEWDN    353
PfGN01_030007300-t41_1-p1      MDENDNRLNNNYRSGFDDTKKNNINDDKGMNFVRDEFYDEYNNNNMNTSPEYMFSQEWDN    353
PfKE01_030006300-t41_1-p1      MDENDNRLNNNYRSGFDDTKKNNINDDKGMNFVRDEFYDEYNNNNMNTSPEYMFSQEWDN    353
PfSN01_030007100-t41_1-p1      MDENDNRLNNNYRSGFDDTKKNNINDDKGMNFVRDEFYDEYNNNNMNTSPEYMFSQEWDN    353
PfTG01_030008300-t41_1-p1      MDENDNRLNNNYRSGFDDTKKNNINDDKGMNFVRDEFYDEYNNNNMNTSPEYMFSQEWDN    353
Pf7G8-2_000071200.1-p1         MDENVNRLNNNYKSGFDDTKKNNINDDKGMNFVRDEFYDEYNNNNMNTSPEYMFSQEWDN    353
Pf7G8_030007400-t41_1-p1       MDENVNRLNNNYKSGFDDTKKNNINDDKGMNFVRDEFYDEYNNNNMNTSPEYMFSQEWDN    353
PfHB3_030005500-t41_1-p1       MDENVNRLNNNYKSGFDDTKKNNINDDKGMNFVRDEFYDEYNNNNMNTSPEYMFSQEWDN    353
PF3D7_0301600.1-p1             MDENDNRLNNNYRSGFDDTKKNNINDDKGMNFVRDEFYDEYNNNNMNTSPEYMFSQEWDN    353
PfNF54_030006800.1-p1          MDENDNRLNNNYRSGFDDTKKNNINDDKGMNFVRDEFYDEYNNNNMNTSPEYMFSQEWDN    353
PfGA01_030008000-t41_1-p1      MDENDNRLNNNYRSGFDDTKKNNINDDKGMNFVRDEFYDEYNNNNMNTSPEYMFSQEWDN    353
PfDd2_030006600-t41_1-p1       MDENDNRLNNNYRSGFDDTKKNNINDDKGMNFVRDEFYDEYNNNNMNTSPEYMFSQEWDN    353
                               **** * *****: ***:: ******* *:***********:* **** *:*:** ****
```

```
PADL01_0034200-t36_1-p1        QNTYDGFESGWNNYNSEYNDFGDQVKVETPYIRVVEEINDESDNIRDDYDMLNNNSSCNR    419
PGABG01_0300800-t36_1-p1       QNTYDGFESGWNNYNSEYNDFVDQVNVETPYIRVVEEINDESDNIRDDYDMLNNDSPCNR    419
PGSY75_0301600-t31_1-p1        QNTYDGFESGWNNYNSEYNDFVDQVNVETPYIRVVEEINDESDNIRDDYDMLNNDSPCNR    419
PBLACG01_0301900-t36_1-p1      QNTYDGFTSGWDNYNSTNSDHRDQVKVETPYIRVVEEIDDESNNIRDDFDILNKDSSCNR    419
PRG01_0304500-t36_1-p1         ENTHDGFSSGWNNYNSENSDHKDQIKVETPYIRVVEEIDDESNNKRDNYDMLNKDSSCNR    419
PRCDC_0300700.1-p1             ENTHDGFSSGWNNYNSENSDHKDQIKVETPYIRVVEEIDDESNNKRDNYDMLNKDSSCNR    419
PfML01_030006900-t41_1-p1      QNTHDGFPSGWNNYNSENSDHKDQIKVETPYIRVVEEVNDESNNKRDNYDTLNKDSSCNR    420
PfGB4_030007500-t41_1-p1       QNTHDGFPSGWNNYNSENSDHKDQIKVETPYIRVVEEVNDESNNKRDNYDTLNKDSSCNR    420
PfSD01_030006800-t41_1-p1      QNTHDGFPSGWNNYNSENSDHKDQIKVETPYIRVVEEVNDESNNKRDNYDTLNKDSSCNR    420
PfKH02_030007200-t41_1-p1      QNTHDGFPSGWNNYNSENSDHKDQIKVETPYIRVVEEVNDESNNKRDNYDTLNKDSSCNR    413
PfIT_030006400-t41_1-p1        QNTHDGFPSGWNNYNSENSDHKDQIKVETPYIRVVEEVNDESNNKRDNYDTLNKDSSCNR    413
PfKH01_030006400-t41_1-p1      QNTHDGFPSGWNNYNSENSDHKDQIKVETPYIRVVEEVNDESNNKRDNYDTLNKDSSCNR    413
PfNF135_030006800.1-p1         QNTHDGFPSGWNNYNSENSDHKDQIKVETPYIRVVEEVDDESNNKRDNYDTLNKDSSCNR    413
PfCD01_030007000-t41_1-p1      QNTHDGFPSGWNNYNSENSDHKDQIKVETPYIRVVEEVNDESNNKRDNYDTLNKDSSCNR    413
PfGN01_030007300-t41_1-p1      QNTHDGFPSGWNNYNSENSDHKDQIKVETPYIRVVEEVNDESNNKRDNYDTLNKDSSCNR    413
PfKE01_030006300-t41_1-p1      QNTHDGFPSGWNNYNSENSDHKDQIKVETPYIRVVEEVNDESNNKRDNYDTLNKDSSCNR    413
PfSN01_030007100-t41_1-p1      QNTHDGFPSGWNNYNSENSDHKDQIKVETPYIRVVEEVNDESNNKRDNYDTLNKDSSCNR    413
PfTG01_030008300-t41_1-p1      QNTHDGFPSGWNNYNSENSDHKDQIKVETPYIRVVEEVNDESNNKRDNYDTLNKDSSCNR    413
Pf7G8-2_000071200.1-p1         QNTHDGFPSGWNNYNSENSDHKDQIKVETPYIRVFEEVNDESNNKRDNYDTLNKDSSCNR    413
Pf7G8_030007400-t41_1-p1       QNTHDGFPSGWNNYNSENSDHKDQIKVETPYIRVFEEVNDESNNKRDNYDTLNKDSSCNR    413
PfHB3_030005500-t41_1-p1       QNTHDGFPSGWNNYNSENSDHKDQIKVETPYIRVVEEVNDESNNKRDNYDTLNKDSSCNR    413
PF3D7_0301600.1-p1             QNTHDGFPSGWNNYNSENSDHKDQIKVETPYIRVVEEVNDESNNKRDNYDTLNKDSSCNR    413
PfNF54_030006800.1-p1          QNTHDGFPSGWNNYNSENSDHKDQIKVETPYIRVVEEVNDESNNKRDNYDTLNKDSSCNR    413
PfGA01_030008000-t41_1-p1      QNTHDGFPSGWNNYNSENSDHKDQIKVETPYIRVVEEVNDESNNKRDNYDTLNKDSSCNR    413
PfDd2_030006600-t41_1-p1       QNTHDGFPSGWNNYNSENSDHKDQIKVETPYIRVVEEVNDESNNKRDNYDTLNKDSSCNR    413
                               :**:*** ***:****  .*. **::********.**::***:* **::* **::* ***

PADL01_0034200-t36_1-p1        NEKMEYYFQSDKFPVDCKGIYTNSNIVVEEYDTPIDVEDTNNNVYSCSDRLNNERQKKHM    479
PGABG01_0300800-t36_1-p1       NEKMEYYFQSDKFPVNCKGIYTNSNIVVEEYDIPIDIKDPNNNVYSYNDRLNNERQKKHM    479
PGSY75_0301600-t31_1-p1        NEKMEYYFQSDKFPVNCKGIYTNSNIVVEEYDIPIDIKDPNNNVYSYNDRLNNERQKKHM    479
PBLACG01_0301900-t36_1-p1      NEKMEYYFQSDKFPVDCKGIYKNSNIVVEEYDTPVDLKEPDNNVYSFNERLNNERQKKYM    479
PRG01_0304500-t36_1-p1         NEKMEYYFQSDKFPVDCKGMYTNSNIVVEEYDTPVDLKETNNNFYSFNEKLNNERPKKYM    479
PRCDC_0300700.1-p1             NEKMEYYFQSDKFPVDCKGMYTNSNIVVEEYDTPVDLKETNNNFYSFNEKLNNERPKKYM    479
PfML01_030006900-t41_1-p1      NEKMEYYFQSDKFPVDCKGIYTNSNIIVEEYDTPVDLKETNDNFYSFNEKLNNERQKKYM    480
PfGB4_030007500-t41_1-p1       NEKMEYYFQSDKFPVDCKGIYTNSNIIVEEYDTPVDLKETNDNFYSFNEKLNNERQKKYM    480
PfSD01_030006800-t41_1-p1      NEKMEYYFQSDKFPVDCKGIYTNSNIIVEEYDTPVDLKETNDNFYSFNEKLNNERQKKYM    480
PfKH02_030007200-t41_1-p1      NEKMEYYFQSDKFPVDCKGIYTNSNIIVEEYDTPVDLKETNDNFYSFNEKLNNERQKKYM    473
PfIT_030006400-t41_1-p1        NEKMEYYFQSDKFPVDCKGIYTNSNIIVEEYDTPVDLKETNDNFYSFNEKLNNERQKKYM    473
PfKH01_030006400-t41_1-p1      NEKMEYYFQSDKFPVDCKGIYTNSNIIVEEYDTPVDLKETNDNFYSFNEKLNNERQKKYM    473
PfNF135_030006800.1-p1         NEKMEYYFQSDKFPVDCKGIYTNSNIIVEEYDTPVDLKETNDNFYSFNEKLNNERQKKYM    473
PfCD01_030007000-t41_1-p1      NEKMEYYFQSDKFPVDCKGIYTNSNIIVEEYDTPVDLKETNDNFYSFNEKLNNERQKKYM    473
PfGN01_030007300-t41_1-p1      NEKMEYYFQSDKFPVDCKGIYTNSNIIVEEYDTPVDLKETNDNFYSFNEKLNNERQKKYM    473
PfKE01_030006300-t41_1-p1      NEKMEYYFQSDKFPVDCKGIYTNSNIIVEEYDTPVDLKETNDNFYSFNEKLNNERQKKYM    473
PfSN01_030007100-t41_1-p1      NEKMEYYFQSDKFPVDCKGIYTNSNIIVEEYDTPVDLKETNDNFYSFNEKLNNERQKKYM    473
PfTG01_030008300-t41_1-p1      NEKMEYYFQSDKFPVDCKGIYTNSNIIVEEYDTPVDLKETNDNFYSFNEKLNNERQKKYM    473
Pf7G8-2_000071200.1-p1         NEKMEYYFQSDKFPVDCKGIYTNSNIIVEEYDTPVDLKETNDNFYSFNEKLNNERQKKYM    473
Pf7G8_030007400-t41_1-p1       NEKMEYYFQSDKFPVDCKGIYTNSNIIVEEYDTPVDLKETNDNFYSFNEKLNNERQKKYM    473
PfHB3_030005500-t41_1-p1       NEKMEYYFQSDKFPVDCKGIYTNSNIIVEEYDTPVDLKETNDNFYSFNEKLNNERQKKYM    473
PF3D7_0301600.1-p1             NEKMEYYFQSDKFPVDCKGIYTNSNIIVEEYDTPVDLKETNDNFYSFNEKLNNERQKKYM    473
PfNF54_030006800.1-p1          NEKMEYYFQSDKFPVDCKGIYTNSNIIVEEYDTPVDLKETNDNFYSFNEKLNNERQKKYM    473
PfGA01_030008000-t41_1-p1      NEKMEYYFQSDKFPVDCKGIYTNSNIIVEEYDTPVDLKETNDNFYSFNEKLNNERQKKYM    473
PfDd2_030006600-t41_1-p1       NEKMEYYFQSDKFPVDCKGIYTNSNIIVEEYDTPVDLKETNDNFYSFNEKLNNERQKKYM    473
                               ***************:***:*.****:***** *:*::: ::*.** .::***** **:*
```

```
PADL01_0034200-t36_1-p1        QEDKEFVNEKFKNDIGHTPSSNSS--------------------------------SSHS    507
PGABG01_0300800-t36_1-p1       QEDKEFVNEKFKNDIGYTPSSNSS--------------------------------SPHS    507
PGSY75_0301600-t31_1-p1        QQDKEFVNEKFKNDIGYTPSSNSS--------------------------------FPHS    507
PBLACG01_0301900-t36_1-p1      QEDKEFKNEKFKNDIGHMSSPSSTLSNSSS------------PHMPSSH----------S    517
PRG01_0304500-t36_1-p1         PEDKEFIN-----EIGHMSSTWSNSSSSHSPSSHFSSSHSSSSHSPSSHSPSSHFSSSHS    534
PRCDC_0300700.1-p1             PEDKEFIN-----EIGHMSSTWSNSSSSHSSSSHSS-----SSHFASSHSSSSHFSS---    526
PfML01_030006900-t41_1-p1      PEDKEFIN-----EMGHMSSTS--------------------------------------    497
PfGB4_030007500-t41_1-p1       PEDKEFIN-----EMGHMSSTS--------------------------------------    497
PfSD01_030006800-t41_1-p1      PEDKEFIN-----EMGHMSSTS--------------------------------------    497
PfKH02_030007200-t41_1-p1      PEDKEFIN-----EMGHMSSTS--------------------------------------    490
PfIT_030006400-t41_1-p1        PEDKEFIN-----EMGHMSSTS--------------------------------------    490
PfKH01_030006400-t41_1-p1      PEDKEFIN-----EMGHMSSTS--------------------------------------    490
PfNF135_030006800.1-p1         PEDKEFIN-----EMGHMSSTS--------------------------------------    490
PfCD01_030007000-t41_1-p1      PEDKEFIN-----EMGHMSSTS--------------------------------------    490
PfGN01_030007300-t41_1-p1      PEDKEFIN-----EMGHMSSTS--------------------------------------    490
PfKE01_030006300-t41_1-p1      PEDKEFIN-----EMGHMSSTS--------------------------------------    490
PfSN01_030007100-t41_1-p1      PEDKEFIN-----EMGHMSSTS--------------------------------------    490
PfTG01_030008300-t41_1-p1      PEDKEFIN-----EMGHMSSTS--------------------------------------    490
Pf7G8-2_000071200.1-p1         PEDKEFIN-----EMGHMSSTS--------------------------------------    490
Pf7G8_030007400-t41_1-p1       PEDKEFIN-----EMGHMSSTS--------------------------------------    490
PfHB3_030005500-t41_1-p1       PEDKEFIN-----EMGHMSSTS--------------------------------------    490
PF3D7_0301600.1-p1             REDKEFIN-----EMGHMSSTS--------------------------------------    490
PfNF54_030006800.1-p1          REDKEFIN-----EMGHMSSTS--------------------------------------    490
PfGA01_030008000-t41_1-p1      PEDKEFIN-----EMGHMSSTS--------------------------------------    490
PfDd2_030006600-t41_1-p1       PEDKEFIN-----EMGHMSSTS--------------------------------------    490
                                :**** *     ::*:  *

PADL01_0034200-t36_1-p1        QPMFTP------------------------------------------ETFQTSNIESPE    525
PGABG01_0300800-t36_1-p1       QPMFTP------------------------------------------ETFQTSNIESPE    525
PGSY75_0301600-t31_1-p1        QPMFTP------------------------------------------ETFQTSNIESPE    525
PBLACG01_0301900-t36_1-p1      QPSFTPETFQTSNIESSEFEEPNIESPEFEEPNIESPEFEEPNI--ESPEFETSNIDSPN    575
PRG01_0304500-t36_1-p1         PPSHSSSSHSPPSHSSSSH-SPPSHSPPSHSSSSHSPPSHSPSSHFPSPH-SQPSFIPET    592
PRCDC_0300700.1-p1             -------SHSPPSHSSSSH-SPPSHSPP-----SHSPPSHSPSSHFPSPH-SQPSFIPET    572
PfML01_030006900-t41_1-p1      --------------------------------------SNSPSSHFPSPH-SQPSFIPET    518
PfGB4_030007500-t41_1-p1       --------------------------------------SNSPSSHFPSPH-SQPSFIPET    518
PfSD01_030006800-t41_1-p1      --------------------------------------SNSPSSHFPSPH-SQPSFIPET    518
PfKH02_030007200-t41_1-p1      --------------------------------------SNSPSSHFPSPH-SQPSFIPET    511
PfIT_030006400-t41_1-p1        --------------------------------------SNSPSSHFPSPH-SQPSFIPET    511
PfKH01_030006400-t41_1-p1      --------------------------------------SNSPSSHFPSPH-SQPSFIPET    511
PfNF135_030006800.1-p1         --------------------------------------SNSPSSHFPSPH-SQPSFIPET    511
PfCD01_030007000-t41_1-p1      --------------------------------------SNSPSSHFPSPH-SQPSFIPET    511
PfGN01_030007300-t41_1-p1      --------------------------------------SNSPSSHFPSPH-SQPSFIPET    511
PfKE01_030006300-t41_1-p1      --------------------------------------SNSPSSHFPSPH-SQPSFIPET    511
PfSN01_030007100-t41_1-p1      --------------------------------------SNSPSSHFPSPH-SQPSFIPET    511
PfTG01_030008300-t41_1-p1      --------------------------------------SNSPSSHFPSPH-SQPSFIPET    511
Pf7G8-2_000071200.1-p1         --------------------------------------SNSPSSHFPSPH-SQPSFIPET    511
Pf7G8_030007400-t41_1-p1       --------------------------------------SNSPSSHFPSPH-SQPSFIPET    511
PfHB3_030005500-t41_1-p1       --------------------------------------SNSPSSHFPSPH-SQPSFIPET    511
PF3D7_0301600.1-p1             --------------------------------------SNSPSSHFPSPH-SQPSFIPET    511
PfNF54_030006800.1-p1          --------------------------------------SNSPSSHFPSPH-SQPSFIPET    511
PfGA01_030008000-t41_1-p1      --------------------------------------SNSPSSHFPSPH-SQPSFIPET    511
PfDd2_030006600-t41_1-p1       --------------------------------------SNSPSSHFPSPH-SQPSFIPET    511
                                                                                  .  .:
```

```
PADL01_0034200-t36_1-p1        FQTP----------TFESPPVNTSDYEIPHFNLSEYDL--------------------P-    554
PGABG01_0300800-t36_1-p1       FQTPTFESPEFQIPTFESPPVNTSDYEIPHFNLSEYDLPHLDPSEYDLPHFDPSEYDLP-    584
PGSY75_0301600-t31_1-p1        FQTPTFESPEFQIPTFESPPVNTSDYEIPHFNLSEYDL--------------------P-    564
PBLACG01_0301900-t36_1-p1      FQAPSFESRAFQLPSFESPPVLTSEYEIPNFDLSEYDLPHLDPSEFKPD-----------    624
PRG01_0304500-t36_1-p1         FQESSIESPEFQLPTFESAPVHISEYELPNFDLLEYDLPHFDPSEFRPEIFDESEFKTEI    652
PRCDC_0300700.1-p1             FQESNVESPEFQLPTFESAPVHISEYELPNFDLLEYDLPHFDPSEFRPEIFDESEFKTEI    632
PfML01_030006900-t41_1-p1      FQDSNIESPEFELPTFESATVHTSEYELPNFDILEYDLPHFDPSEFKPEIFDESEL----    574
PfGB4_030007500-t41_1-p1       FQDSNIESPEFELPTFESATVHTSEYELPNFDILEYDLPHFDPSEFKPEIF---------    569
PfSD01_030006800-t41_1-p1      FQDSNIESPEFELPTFESATVHTSEYELPNFDILEYDLPHFDPSEFKPEIF---------    569
PfKH02_030007200-t41_1-p1      FQDSNIESPEFELPTFESATVHTSEYELPNFDILEYDLPHFDPSEFKPEIFDESEF----    567
PfIT_030006400-t41_1-p1        FQDSNIESPEFELPTFESATVHTSEYELPNFDILEYDLPHFDPSEFKPEIFD--------    563
PfKH01_030006400-t41_1-p1      FQDSNIESPEFELPTFESATVHTSEYELPNFDILEYDLPHFDPSEFKPEIFD--------    563
PfNF135_030006800.1-p1         FQDSNIESPEFELPTFESATVHTSEYELPNFDILEYDLPHFDPSEFKPEIFD--------    563
PfCD01_030007000-t41_1-p1      FQDSNIESPEFELPTFESATVHTSEYELPNFDILEYDLPHFDPSEFKPEIFD--------    563
PfGN01_030007300-t41_1-p1      FQDSNIESPEFELPTFESATVHTSEYELPNFDILEYDLPHFDPSEFKPEIFD--------    563
PfKE01_030006300-t41_1-p1      FQDSNIESPEFELPTFESATVHTSEYELPNFDILEYDLPHFDPSEFKPEIFD--------    563
PfSN01_030007100-t41_1-p1      FQDSNIESPEFELPTFESATVHTSEYELPNFDILEYDLPHFDPSEFKPEIFD--------    563
PfTG01_030008300-t41_1-p1      FQDSNIESPEFELPTFESATVHTSEYELPNFDILEYDLPHFDPSEFKPEIFD--------    563
Pf7G8-2_000071200.1-p1         FQDSNIESPEFELPTFESATVHTSEYELPNFDILEYDLPHFDPSEFKPEIFD--------    563
Pf7G8_030007400-t41_1-p1       FQDSNIESPEFELPTFESATVHTSEYELPNFDILEYDLPHFDPSEFKPEIFD--------    563
PfHB3_030005500-t41_1-p1       FQDSNIESPEFELPTFESATVHTSEYELPNFDILEYDLPHFDPSEFKPEIFD--------    563
PF3D7_0301600.1-p1             FQDSNIESPEFELPTFESATVHTSEYELPNFDILEYDLPHFDPSEFKPEIFDESEF----    567
PfNF54_030006800.1-p1          FQDSNIESPEFELPTFESATVHTSEYELPNFDILEYDLPHFDPSEFKPEIFDESEF----    567
PfGA01_030008000-t41_1-p1      FQDSNIESPEFELPTFESATVHTSEYELPNFDILEYDLPHFDPSEFKPEIFDESEF----    567
PfDd2_030006600-t41_1-p1       FQDSNIESPEFELPTFESATVHTSEYELPNFDILEYDLPHFDPSEFKPEIFDESEF----    567
                               **            :***  *  *:**:*:*:: ****

PADL01_0034200-t36_1-p1        ---------HLDPSEFKPEPFDESEFDPPLFDPSEYKNSSFKQQKYSASSNSSRSYENEF    605
PGABG01_0300800-t36_1-p1       ---------HLDPSEFKPEPFDESEFDPPVFDPSEYKNSTFKQPKYSTSSHSSRSYENEF    635
PGSY75_0301600-t31_1-p1        ---------HLDPSEFKPEPFDESEFDPPVFDPSEYKNSTFKQPKYSTSSHSSRSYENEF    615
PBLACG01_0301900-t36_1-p1      -------------------PFDESDFKPQVFDPSEYKNSTLKQSKLSTSSNSAYSYENEF    665
PRG01_0304500-t36_1-p1         FDESEF----------RPEIFDESEFKPSVFDTSEYKNSTFKESKFSTSSHSTRSYENEF    702
PRCDC_0300700.1-p1             FDESEFRPEIFDESEFRPEIFDESEFKPSVFDPSEYKNSTFKESKFSTSSHSTRSYENEF    692
PfML01_030006900-t41_1-p1      ------K----------PEIFDESEFKPSVFDPSEYKNSPFKESKISTSSHSTRSYENEF    618
PfGB4_030007500-t41_1-p1       ---------------------DESEFKPSVFDPSEYKNSPFKESKISTSSHSTRSYENEF    608
PfSD01_030006800-t41_1-p1      ---------------------DESEFKPSVFDPSEYKNSPFKESKISTSSHSTRSYENEF    608
PfKH02_030007200-t41_1-p1      ------KPEIFDESEFKPEIFDESEFKPSVFDPSEYKNSPFKESKISTSSHSTRSYENEF    621
PfIT_030006400-t41_1-p1        ----------------------ESEFKPSVFDPSEYKNSPFKESKISTSSHSTRSYENEF    601
PfKH01_030006400-t41_1-p1      ----------------------ESEFKPSVFDPSEYKNSPFKESKISTSSHSTRSYENEF    601
PfNF135_030006800.1-p1         ----------------------ESEFKPSVFDPSEYKNSPFKESKISTSSHSTRSYENEF    601
PfCD01_030007000-t41_1-p1      ----------------------ESEFKPSVFDPSEYKNSPFKESKISTSSHSTRSYENEF    601
PfGN01_030007300-t41_1-p1      ----------------------ESEFKPSVFDPSEYKNSPFKESKISTSSHSTRSYENEF    601
PfKE01_030006300-t41_1-p1      ----------------------ESEFKPSVFDPSEYKNSPFKESKISTSSHSTRSYENEF    601
PfSN01_030007100-t41_1-p1      ----------------------ESEFKPSVFDPSEYKNSPFKESKISTSSHSTRSYENEF    601
PfTG01_030008300-t41_1-p1      ----------------------ESEFKPSVFDPSEYKNSPFKESKISTSSHSTRSYENEF    601
Pf7G8-2_000071200.1-p1         ----------------------ESEFKPSVFDPSEYKNSPFKESKISTSSHSTRSYENEF    601
Pf7G8_030007400-t41_1-p1       ----------------------ESEFKPSVFDPSEYKNSPFKESKISTSSHSTRSYENEF    601
PfHB3_030005500-t41_1-p1       ----------------------ESEFKPSVFDPSEYKNSPFKESKISTSSHSTRSYENEF    601
PF3D7_0301600.1-p1             ------K----------PEIFDESEFKPSVFDPSEYKNSPFKESKISTSSHSTRSYENEF    611
PfNF54_030006800.1-p1          ------K----------PEIFDESEFKPSVFDPSEYKNSPFKESKISTSSHSTRSYENEF    611
PfGA01_030008000-t41_1-p1      ------K----------PEIFDESEFKPSVFDPSEYKNSPFKESKISTSSHSTRSYENEF    611
PfDd2_030006600-t41_1-p1       ------K----------PEIFDESEFKPSVFDPSEYKNSPFKESKISTSSHSTRSYENEF    611
                                                     **:*.* :** ****** :*: * *:**:*: ******
```

```
PADL01_0034200-t36_1-p1         NLSTHRRNCESSSRIHVDDENYKRLKNILSSLCSSREKIYDLSKEEKEFLVKMLKLDYED    665
PGABG01_0300800-t36_1-p1        NLSTHRRNCESSSRIHVDDENYMRLKNILSSLCSSREKIYDLSKEEKEFLVKMLKLDYED    695
PGSY75_0301600-t31_1-p1         NLSTHRRNCESSSRIHVDDENYMRLKNILSSLCSSREKIYDLSKEEKEFLVKMLKLDYED    675
PBLACG01_0301900-t36_1-p1       NLSTHRRNSESSSRIHVDDENYKRLKNILSNLCSSREKIYDLSKEEKEFLVKMLKLDYED    725
PRG01_0304500-t36_1-p1          NLSSDRRNSESSSRIHVDDENYKRLKDILSNLCSSREKIYDLSKEEKEFLVKMLKLDYED    762
PRCDC_0300700.1-p1              NLSSDRRNSESSSRIHVDDENYKRLKDILSNLCSSREKIYDLSKEEKEFLVKMLKLDYED    752
PfML01_030006900-t41_1-p1       NLSTDRRNSESSSRIHVDDENYKRLKKILSNLCSSREKIYDLSKEEKEFLVKMLQLDYED    678
PfGB4_030007500-t41_1-p1        NLSTDRRNSESSSRIHVDDENYKRLKKILSNLCSSREKIYDLSKEEKEFLVKMLQLDYED    668
PfSD01_030006800-t41_1-p1       NLSTDRRNSESSSRIHVDDENYKRLKKILSNLCSSREKIYDLSKEEKEFLVKMLQLDYED    668
PfKH02_030007200-t41_1-p1       NLSTDRRNSESSSRIHVDDENYKRLKKILSNLCSSREKIYDLSKEEKQFLVKMLQLDYED    681
PfIT_030006400-t41_1-p1         NLSTDRRNSESSSRIHVDDENYKRLKKILSNLCSSREKIYDLSKEEKEFLVKMLQLDYED    661
PfKH01_030006400-t41_1-p1       NLSTDRRNSESSSRIHVDDENYKRLKKILSNLCSSREKIYDLSKEEKQFLVKMLQLDYED    661
PfNF135_030006800.1-p1          NLSTDRRNSESSSRIHVDDENYKRLKKILSNLCSSREKIYDLSKEEKEFLVKMLQLDYED    661
PfCD01_030007000-t41_1-p1       NLSTDRRNSESSSRIHVDDENYKRLKKILSNLCSSREKIYDLSKEEKEFLVKMLQLDYED    661
PfGN01_030007300-t41_1-p1       NLSTDRRNSESSSRIHVDDENYKRLKKILSNLCSSREKIYDLSKEEKEFLVKMLQLDYED    661
PfKE01_030006300-t41_1-p1       NLSTDRRNSESSSRIHVDDENYKRLKKILSNLCSSREKIYDLSKEEKEFLVKMLQLDYED    661
PfSN01_030007100-t41_1-p1       NLSTDRRNSESSSRIHVDDENYKRLKKILSNLCSSREKIYDLSKEEKEFLVKMLQLDYED    661
PfTG01_030008300-t41_1-p1       NLSTDRRNSESSSRIHVDDENYKRLKKILSNLCSSREKIYDLSKEEKEFLVKMLQLDYED    661
Pf7G8-2_000071200.1-p1          NLSTDRRNSESSSRIHVDDENYKRLKKILSNLCSSREKIYDLSKEEKEFLVKMLQLDYED    661
Pf7G8_030007400-t41_1-p1        NLSTDRRNSESSSRIHVDDENYKRLKKILSNLCSSREKIYDLSKEEKEFLVKMLQLDYED    661
PfHB3_030005500-t41_1-p1        NLSTDRRNSESSSRIHVDDENYKRLKKILSNLCSSREKIYDLSKEEKEFLVKMLQLDYED    661
PF3D7_0301600.1-p1              NLSTDRRNSESSSRIHVDDENYKRLKKILSNLCSSREKIYDLSKEEKEFLVKMLQLDYED    671
PfNF54_030006800.1-p1           NLSTDRRNSESSSRIHVDDENYKRLKKILSNLCSSREKIYDLSKEEKEFLVKMLQLDYED    671
PfGA01_030008000-t41_1-p1       NLSTDRRNSESSSKIHVDDENYKRLKKILSNLCSSREKIYDLSKEEKEFLVKMLQLDYED    671
PfDd2_030006600-t41_1-p1        NLSTDRRNSESSSRIHVDDENYKRLKKILSNLCSSREKIYDLSKEEKEFLVKMLQLDYED    671
                                ***:.***.****:******** ***.***.****************:******:*****

PADL01_0034200-t36_1-p1         FNFNYRENPRIYNESSSFTNFELKRIHSKIKAFIYKCLLRLLQSYEKSTENHGCANIYDI    725
PGABG01_0300800-t36_1-p1        FNFNSRQNPRIYNESTSFTNFELKRIHSKIKAFIYKCLLRLLQSYEKSSENHGYANIYDI    755
PGSY75_0301600-t31_1-p1         FNFNSRQNARIYNESSSFTNFELKRIHSKIKAFIYKCLLRLLQSYEKSSENHGYANIYDI    735
PBLACG01_0301900-t36_1-p1       FNFNYRGNTRHYNESSPFTNFELKRIHLKVKAFIYKCLLRLLHSCEKSTGNYVDSNIYDI    785
PRG01_0304500-t36_1-p1          FNFNSRGNTRNYNDSSPFTNFELKRIHLKIKAFIYKCLLRLLQSYEKTTENHVGSNIYDI    822
PRCDC_0300700.1-p1              FNFNSRGNTRNYNDSSPFTNFELKRIHLKIKAFIYKCLLRLLQSYEKTTENHVGSNIYDI    812
PfML01_030006900-t41_1-p1       FNFNYRGNTRSYNDSSPFTNFELKRIHLKIKAFIYKCLLRLLQSYEKSTESRVGSNIYDI    738
PfGB4_030007500-t41_1-p1        FNFNYRGNTRSYNDSSPFTNFELKRIHLKIKAFIYKCLLRLLQSYEKSTESRVGSNIYDI    728
PfSD01_030006800-t41_1-p1       FNFNYRGNTRSYNDSSPFTNFELKRIHLKIKEFIYKCLLRLLQSYEKSTESRVGSNIYDI    728
PfKH02_030007200-t41_1-p1       FNFNYRGNTRSYNDSSPFTNFELKRIHLKIKAFIYKCLLRLLQSYEKSTESRVGSNIYDI    741
PfIT_030006400-t41_1-p1         FNFNYRGNTRSYNDSSPFTNFELKRIHLKIKAFIYKCLLRLLQSYEKSTESRVGSNIYDI    721
PfKH01_030006400-t41_1-p1       FNFNYRGNTRSYNDSSPFTNFELKRIHLKIKAFIYKCLLRLLQSYEKSTESRVGSNIYDI    721
PfNF135_030006800.1-p1          FNFNYRGNTRSYNDSSPFTNFELKRIHLKIKAFIYKCLLRLLQSYEKSTESRVGSNIYDI    721
PfCD01_030007000-t41_1-p1       FNFNYRGNTRSYNDSSPFTNFELKRIHLKIKAFIYKCLLRLLQSYEKSTESRVGSNIYDI    721
PfGN01_030007300-t41_1-p1       FNFNYRGNTRSYNDSSPFTNFELKRIHLKIKAFIYKCLLRLLQSYEKSTESRVGSNIYDI    721
PfKE01_030006300-t41_1-p1       FNFNYRGNTRSYNDSSPFTNFELKRIHLKIKAFIYKCLLRLLQSYEKSTESRVGSNIYDI    721
PfSN01_030007100-t41_1-p1       FNFNYRGNTRSYNDSSPFTNFELKRIHLKIKAFIYKCLLRLLQSYEKSTESRVGSNIYDI    721
PfTG01_030008300-t41_1-p1       FNFNYRGNTRSYNDSSPFTNFELKRIHLKIKAFIYKCLLRLLQSYEKSTESRVGSNIYDI    721
Pf7G8-2_000071200.1-p1          FNFNYRGNTRSYNDSSPFTNFELKRIHLKIKAFIYKCLLRLLQSYEKSTESRVGSNIYDI    721
Pf7G8_030007400-t41_1-p1        FNFNYRGNTRSYNDSSPFTNFELKRIHLKIKAFIYKCLLRLLQSYEKSTESRVGSNIYDI    721
PfHB3_030005500-t41_1-p1        FNFNYRGNTRSYNDSSPFTNFELKRIHLKIKAFIYKCLLRLLQSYEKSTESRVGSNIYDI    721
PF3D7_0301600.1-p1              FNFNYRGNTRSYNDSSPFTNFELKRIHLKIKAFIYKCLLRLLQSYEKSTESRVGSNIYDI    731
PfNF54_030006800.1-p1           FNFNYRGNTRSYNDSSPFTNFELKRIHLKIKAFIYKCLLRLLQSYEKSTESRVGSNIYDI    731
PfGA01_030008000-t41_1-p1       FNFNYRGNTRSYNDSSPFTNFELKRIHLKIKAFIYKCLLRLLQSYEKSTESRVGSNIYDI    731
PfDd2_030006600-t41_1-p1        FNFNYRGNTRSYNDSSPFTNFELKRIHLKIKAFIYKCLLRLLQSYEKSTESRVGSNIYDI    731
                                **** * * * **:*: ********** *:* **********:* **:: .   :*****
```

| | | |
|---|---|---|
| PADL01_0034200-t36_1-p1 | Y | 726 |
| PGABG01_0300800-t36_1-p1 | Y | 756 |
| PGSY75_0301600-t31_1-p1 | Y | 736 |
| PBLACG01_0301900-t36_1-p1 | Y | 786 |
| PRG01_0304500-t36_1-p1 | Y | 823 |
| PRCDC_0300700.1-p1 | Y | 813 |
| PfML01_030006900-t41_1-p1 | Y | 739 |
| PfGB4_030007500-t41_1-p1 | Y | 729 |
| PfSD01_030006800-t41_1-p1 | Y | 729 |
| PfKH02_030007200-t41_1-p1 | Y | 742 |
| PfIT_030006400-t41_1-p1 | Y | 722 |
| PfKH01_030006400-t41_1-p1 | Y | 722 |
| PfNF135_030006800.1-p1 | Y | 722 |
| PfCD01_030007000-t41_1-p1 | Y | 722 |
| PfGN01_030007300-t41_1-p1 | Y | 722 |
| PfKE01_030006300-t41_1-p1 | Y | 722 |
| PfSN01_030007100-t41_1-p1 | Y | 722 |
| PfTG01_030008300-t41_1-p1 | Y | 722 |
| Pf7G8-2_000071200.1-p1 | Y | 722 |
| Pf7G8_030007400-t41_1-p1 | Y | 722 |
| PfHB3_030005500-t41_1-p1 | Y | 722 |
| PF3D7_0301600.1-p1 | Y | 732 |
| PfNF54_030006800.1-p1 | Y | 732 |
| PfGA01_030008000-t41_1-p1 | Y | 732 |
| PfDd2_030006600-t41_1-p1 | Y | 732 |
| | * | |

**Alignment Syntenic Orthologues PF3D7_0113300**

```
PADL01_0110900-t36_1-p1        ------MKNIKRELFFFFVLLFFLYIICGCIYENFHECHYNKIFNNPHASEKYRQFNKIR    54
PGABG01_0111700-t36_1-p1       ------MKNIKRELFLFFVLLFFLYIICGCIYEKLYECQYNKIFNNPHESEKYRQFNKIR    54
PGSY75_0113300-t31_1-p1        ------MKNIKRELFLFFVVLFFLYIICGCIYEKLYECQYNKIYNNPHESEKYRQFNKIR    54
SPJ08214.1                     ------MKNIKSEGLIFFLLLFFIYIIFGCIHENLHECSYKKTLKSLHVSEKYREFNKIR    54
PF3D7_0113300.1-p1             MKNIKNMKNIKSEGFIFFVLVFFICIIFGCIYESLHEGPYKKTLNSLHESTKYRHFNKIR    60
PfNF54_010017900.1-p1          MKNIKNMKNIKSEGFIFFVLVFFICIIFGCIYESLHEGPYKKTLNSLHESTKYRHFNKIR    60
PfGB4_010017100-t41_1-p1       MKNIKNMKNIKSEGFIFFVLVFFICIIFGCIYESLHEGPYKKTLNSLHESTKYRHFNKIR    60
SOS76152.1                     MKNIKNMKNIKSEGFIFFVLVFFICIIFGCIYESLHECPYKKTLNSLHESTKYRHFNKIR    60
PfHB3_010017200-t41_1-p1       MKNIKNMKNIKSEGFIFFVLVFFICIIFGCIYESLHEGPYKKTLNSLHESTKYRHFNKIR    60
PfTG01_010018100-t41_1-p1      MKNIKNMKNIKSEGFIFFVLVFFICIIFGCIYESLHEGPYKKTLNSLHESTKYRHFNKIR    60
PfNF135_010017200.1-p1         MKNIKNMKNIKSEGFIFFVLVFFICIIFGCIYESLHEGPYKKTLNSLHESTKYRHFNKIR    60
PfKH02_010016900-t41_1-p1      MKNIKNMKNIKSEGFIFFVLVFFICIIFGCIYESLHEGPYKKTLNSLHESTKYRHFNKIR    60
PfNF166_010016800.1-p1         MKNIKNMKNIKSEGFIFFVLVFFICIIFGCIYESLHEGPYKKTLNSLHESTKYRHFNKIR    60
Pf7G8-2_000046900.1-p1         MKNIKNMKNIKSEGFIFFVLVFFICIIFGCIYESLHEGPYKKTLNSLHESTKYRHFNKIR    60
Pf7G8_010017300-t41_1-p1       MKNIKNMKNIKSEGFIFFVLVFFICIIFGCIYESLHEGPYKKTLNSLHESTKYRHFNKIR    60
PfKE01_010016300-t41_1-p1      MKNIKNMKNIKSEGFIFFVLVFFICIIFGCIYESLHEGPYKKTLNSLHESTKYRHFNKIR    60
PfIT_010016400-t41_1-p1        MKNIKNMKNIKSEGFIFFVLVFFICIIFGCIYESLHEGPYKKTLNSLHESTKYRHFNKIR    60
PfSN01_010015800-t41_1-p1      MKNIKNMKNIKSEGFIFFVLVFFICIIFGCIYESLHEGPYKKTLNSLHESTKYRHFNKIR    60
PfCD01_010017400-t41_1-p1      MKNIKNMKNIKSEGFIFFVLVFFICIIFGCIYESLHEGPYKKTLNSLHESTKYRHFNKIR    60
PfSD01_010016600-t41_1-p1      MKNIKNMKNIKSEGFIFFVLVFFICIIFGCIYESLHEGPYKKTLNSLHESTKYRHFNKIR    60
PfKH01_010018100-t41_1-p1      MKNIKNMKNIKSEGFIFFVLVFFICIIFGCIYESLHEGPYKKTLNSLHESTKYRHFNKIR    60
PfGA01_010017200-t41_1-p1      MKNIKNMKNIKSEGFIFFVLVFFICIIFGCIYESLHEGPYKKTLNSLHESTKYRHFNKIR    60
PfGN01_010017400-t41_1-p1      MKNIKNMKNIKSEGFIFFVLVFFICIIFGCIYESLHEGPYKKTLNSLHESTKYRHFNKIR    60
PfDd2_010016800-t41_1-p1       MKNIKNMKNIKSEGFIFFVLVFFICIIFGCIYESLHEGPYKKTLNSLHESTKYRHFNKIR    60
PfML01_010016900-t41_1-p1      MKNIKNMKNIKSEGFIFFVLVFFICIIFGCIYESLHEGPYKKTLNSLHESTKYRHFNKIR    60
PRCDC_0111600.1-p1             ------MKNIKSEGFIFFVLVFFICIIFGCIYESLHEYPYKKALNCLHESEKFREFNKIR    54
PRG01_0113700-t36_1-p1         ------MKNIKSEGFIFFVLVFFICIIFGCIYESLHEYPYKKALNCLHESEKFREFNKIR    54
                                     ***** * ::**:::**: ** ***:*.::*  *:*  :  * * *:*.*****

PADL01_0110900-t36_1-p1        LLSENKNILQFNKEKISLHDYINNDNYNTVDANDYNCYKDKEEQCNDSSC-VDKKNYNRS    113
PGABG01_0111700-t36_1-p1       LLSENKNILEFNKEKISLHDYINNDNYNTLDAIDYNFYNDKEEQCNDTTC-VDKKNYNRT    113
PGSY75_0113300-t31_1-p1        LLSENKNILEFNKEKISLHDYINNDNYNTLDAIDYNFYNDKEEQCNDTTC-VDKKNYNRT    113
SPJ08214.1                     LLTEYKNTLQLKEEKLSLHDYINNDRYNNIDTNDHTLCKDTEEQFNDTTS-VDKKNEKVT    113
PF3D7_0113300.1-p1             LLTEYKDTLQIKVEQKSLRDYVNNDRYNNVNTNDYTSYKDKGEQFNDTICVVDKKKENVT    120
PfNF54_010017900.1-p1          LLTEYKDTLQIKVEQKSLRDYVNNDRYNNVNTNDYTSYKDKGEQFNDTICVVDKKKENVT    120
PfGB4_010017100-t41_1-p1       LLTEYKDTLQIKVEQKSLRDYVNNDRYNNVNTNDYTSYKDKGEQFNDTICVVDKKKENVT    120
SOS76152.1                     LLTEYKDTLQIKVEQKSLRDYVNNDRYNNVNTNDYTSYNDKGEQFNDTICVVDKKKENVT    120
PfHB3_010017200-t41_1-p1       LLTEYKDTLQIKVEQKSLRDYVNNDRYNNVNTNDYTSYKDKGEQFNDTICVVDKKKENVT    120
PfTG01_010018100-t41_1-p1      LLTEYKDTLQIKVEQKSLRDYVNNDRYNNVNTNDYTSYKDKGEQFNDTICVVDKKKENVT    120
PfNF135_010017200.1-p1         LLTEYKDTLQIKVEQKSLRDYVNNDRYNNVNTNDYTSYKDKGEQFNDTICVVDKKKENVT    120
PfKH02_010016900-t41_1-p1      LLTEYKDTLQIKVEQKSLRDYVNNDRYNNVNTNDYTSYKDKGEQFNDTICVVDKKKENVT    120
PfNF166_010016800.1-p1         LLTEYKDTLQIKVEQKSLRDYVNNDRYNNVNTNDYTSYKDKGEQFNDTICVVDKKKENVT    120
Pf7G8-2_000046900.1-p1         LLTEYKDTLQIKVEQKSLRDYVNNDRYNNVNTNDYTSYKDKGEQFNDTICVVDKKKENVT    120
Pf7G8_010017300-t41_1-p1       LLTEYKDTLQIKVEQKSLRDYVNNDRYNNVNTNDYTSYKDKGEQFNDTICVVDKKKENVT    120
PfKE01_010016300-t41_1-p1      LLTEYKDTLQIKVEQKSLRDYVNNDRYNNVNTNDYTSYKDKGEQFNDTICVVDKKKENVT    120
PfIT_010016400-t41_1-p1        LLTEYKDTLQIKVEQKSLRDYVNNDRYNNVNTNDYTSYKDKGEQFNDTICVVDKKKENVT    120
PfSN01_010015800-t41_1-p1      LLTEYKDTLQIKVEQKSLRDYVNNDRYNNVNTNDYTSYKDKGEQFNDTICVVDKKKENVT    120
PfCD01_010017400-t41_1-p1      LLTEYKDTLQIKVEQKSLRDYVNNDRYNNVNTNDYTSYKDKGEQFNDTICVVDKKKENVT    120
PfSD01_010016600-t41_1-p1      LLTEYKDTLQIKVEQKSLRDYVNNDRYNNVNTNDYTSYKDKGEQFNDTICVVDKKKENVT    120
PfKH01_010018100-t41_1-p1      LLTEYKDTLQIKVEQKSLRDYVNNDRYNNVNTNDYTSYKDKGEQFNDTICVVDKKKENVT    120
PfGA01_010017200-t41_1-p1      LLTEYKDTLQIKVEQKSLRDYVNNDRYNNVNTNDYTSYKDKGEQFNDTICVVDKKKENVT    120
PfGN01_010017400-t41_1-p1      LLTEYKDTLQIKVEQKSLRDYVNNDRYNNVNTNDYTSYKDKGEQFNDTICVVDKKKENVT    120
PfDd2_010016800-t41_1-p1       LLTEYKDTLQIKVEQKSLRDYVNNDRYNNVNTNDYTSYKDKGEQFNDTICVVDKKKENVT    120
PfML01_010016900-t41_1-p1      LLTEYKDTLQIKVEQKSLRDYVNNDRYNNVNTNDYTSYKDKGEQFNDTICVVDKKKENVT    120
PRCDC_0111600.1-p1             LLTEYKDTLQIKEEKFTLHDYVNNDRYNNVNTNNYTLYKDKEEQFNDTICVVDKKKENGT    114
PRG01_0113700-t36_1-p1         LLTEYKDTLQIKEEKFTLHDYVNNDRYNNVNTNNYTLYKDKEEQFNDTICVVDKKKENGT    114
                               **:* *: *::: *: :*:**:***.**.::: ::.  :*. ** **: . ****: : :
```

```
PADL01_0110900-t36_1-p1         NNNEEECSKNFYQYLKYMEQNVNVNTNNKNINDKNINDNNTELQIISNDHNTKNYKYDDN    173
PGABG01_0111700-t36_1-p1        NNNEEECSKNFYQYLKYMEQNVNVNTNN----------INTELQIISNDHNKQNYKYDNN    163
PGSY75_0113300-t31_1-p1         NNNEEECSKNFYQYLKYMEQNVNVNTNN----------INTELQIISNDHNKQNYNYDNN    163
SPJ08214.1                      TNNDEKCNKNFYQYLQYLEHNVNGYTNNKNTEL----------QIVSND-----------    152
PF3D7_0113300.1-p1              INNEEECNKNFYQYLQYLEHNNK-------------------------------------    143
PfNF54_010017900.1-p1           INNEEECNKNFYQYLQYLEHNNK-------------------------------------    143
PfGB4_010017100-t41_1-p1        INNEEECNKNFYQYLQYLEHNNK-------------------------------------    143
SOS76152.1                      INNEEECNKNFYQYLQYLEHNNK-------------------------------------    143
PfHB3_010017200-t41_1-p1        INNEEECNKNFYQYLQYLEHNNK-------------------------------------    143
PfTG01_010018100-t41_1-p1       INNEEECNKNFYQYLQYLEHNNK-------------------------------------    143
PfNF135_010017200.1-p1          INNEEECNKNFYQYLQYLEHNNK-------------------------------------    143
PfKH02_010016900-t41_1-p1       INNEEECNKNFYQYLQYLEHNNK-------------------------------------    143
PfNF166_010016800.1-p1          INNEEECNKNFYQYLQYLEHNNK-------------------------------------    143
Pf7G8-2_000046900.1-p1          INNEEECNKNFYQYLQYLEHNNK-------------------------------------    143
Pf7G8_010017300-t41_1-p1        INNEEECNKNFYQYLQYLEHNNK-------------------------------------    143
PfKE01_010016300-t41_1-p1       INNEEECNKNFYQYLQYLEHNNK-------------------------------------    143
PfIT_010016400-t41_1-p1         INNEEECNKNFYQYLQYLEHNNK-------------------------------------    143
PfSN01_010015800-t41_1-p1       INNEEECNKNFYQYLQYLEHNNK-------------------------------------    143
PfCD01_010017400-t41_1-p1       INNEEECNKNFYQYLQYLEHNNK-------------------------------------    143
PfSD01_010016600-t41_1-p1       INNEEECNKNFYQYLQYLEHNNK-------------------------------------    143
PfKH01_010018100-t41_1-p1       INNEEECNKNFYQYLQYLEHNNK-------------------------------------    143
PfGA01_010017200-t41_1-p1       INNEEECNKNFYQYLQYLEHNNK-------------------------------------    143
PfGN01_010017400-t41_1-p1       INNEEECNKNFYQYLQYLEHNNK-------------------------------------    143
PfDd2_010016800-t41_1-p1        INNEEECNKNFYQYLQYLEHNNK-------------------------------------    143
PfML01_010016900-t41_1-p1       INNEEECNKNFYQYLQYLEHNNK-------------------------------------    143
PRCDC_0111600.1-p1              INNEEECNKNFYQYLQYLEHNVHGKENNNKYDN----------KYDNQNDNKYDSKY---    161
PRG01_0113700-t36_1-p1          INNEEECNKNFYQYLQYLEHNVHGKENNNKYDN----------KYDNQNDNKYDSKYDNQ    164
                                 **:*:*.*******:*:*:* :

PADL01_0110900-t36_1-p1         NN-------------------------------SNNNNSEYGKTNNFLERNNKVIDSQQN    202
PGABG01_0111700-t36_1-p1        SDN-----------------------NN-NSDNNNNNNSEYGETNYFLERNNKLIDSQQN    199
PGSY75_0113300-t31_1-p1         SDN-----------------------NNNNSDNNNNNNSEYGETNYFLERNNKLIDSQQN    200
SPJ08214.1                      -----------------------QNIQYY----ENENKNENDQTNYFLGGNNKHMDSQQN    185
PF3D7_0113300.1-p1              ------------------------------------QDNKYEETNYFLQGNDKHIDSEHN    167
PfNF54_010017900.1-p1           ------------------------------------QDNKYEETNYFLQGNDKHIDSEHN    167
PfGB4_010017100-t41_1-p1        ------------------------------------QDNKYEETNYFLQGNDKHIDSEHN    167
SOS76152.1                      ------------------------------------QDNKYEETNYFLQGNDKHIDSEHN    167
PfHB3_010017200-t41_1-p1        ------------------------------------QDNKYEETNYFLQGNDKHIDSEHN    167
PfTG01_010018100-t41_1-p1       ------------------------------------QDNKYEETNYFLQGNDKHIDSEHN    167
PfNF135_010017200.1-p1          ------------------------------------QDNKYEETNYFLQGNDKHIDSEHN    167
PfKH02_010016900-t41_1-p1       ------------------------------------QDNKYEETNYFLQGNDKHIDSEHN    167
PfNF166_010016800.1-p1          ------------------------------------QDNKYEETNYFLQGNDKHIDSEHN    167
Pf7G8-2_000046900.1-p1          ------------------------------------QDNKYEETNYFLQGNDKHIDSEHN    167
Pf7G8_010017300-t41_1-p1        ------------------------------------QDNKYEETNYFLQGNDKHIDSEHN    167
PfKE01_010016300-t41_1-p1       ------------------------------------QDNKYEETNYFLQGNDKHIDSEHN    167
PfIT_010016400-t41_1-p1         ------------------------------------QDNKYEETNYFLQGNDKHIDSEHN    167
PfSN01_010015800-t41_1-p1       ------------------------------------QDNKYEETNYFLQGNDKHIDSEHN    167
PfCD01_010017400-t41_1-p1       ------------------------------------QDNKYEETNYFLQGNDKHIDSEHN    167
PfSD01_010016600-t41_1-p1       ------------------------------------QDNKYEETNYFLQGNDKHIDSEHN    167
PfKH01_010018100-t41_1-p1       ------------------------------------QDNKYEETNYFLQGNDKHIDSEHN    167
PfGA01_010017200-t41_1-p1       ------------------------------------QDNKYEETNYFLQGNDKHIDSEHN    167
PfGN01_010017400-t41_1-p1       ------------------------------------QDNKYEETNYFLQGNDKHIDSEHN    167
PfDd2_010016800-t41_1-p1        ------------------------------------QDNKYEETNYFLQGNDKHIDSEHN    167
PfML01_010016900-t41_1-p1       ------------------------------------QDNKYEETNYFLQGNDKHIDSEHN    167
PRCDC_0111600.1-p1              -DNQNDNKYDSKYDNQYDNKYDNQNDNKYDNQNDNQYDNKYDETNYFLWGNDKHIDSEHN    220
PRG01_0113700-t36_1-p1          YDNQNDNKYDSKYDNQYDNKYDNQNDNKYDNQNDNQYDNKYDETNYFLWGNDKHIDSEHN    224
                                                                     ..:  :** **  *:* :**::*
```

```
PADL01_0110900-t36_1-p1          KMNNIFKETINKTLTYDMSTDNSHTNNNSRHYENQNGNREYMNLSNDNLTYGNSPYDILP    262
PGABG01_0111700-t36_1-p1         EMNNIFQETINKTLTYDMSTENSYTNNNSMHYETQNGNREYMNLSNNNLTYIISPYDILP    259
PGSY75_0113300-t31_1-p1          EMNNIFQETINKTLTYDMSTENSHTNNNSIHYETQNGNREYMNLSNNNLTYIISPYDILP    260
SPJ08214.1                       GINKIYKETIHNTLTYNESTENTHTHNNLRDDKPQNGKMEYNNQSNNNLPYDNSPYNISP    245
PF3D7_0113300.1-p1               GINKMYKETIHKTLTSDVSTENSYTHNNSRDDEPQNGKRTYNNQSNNNLPYDNSSYNISP    227
PfNF54_010017900.1-p1            GINKMYKETIHKTLTSDVSTENSYTHNNSRDDEPQNGKRTYNNQSNNNLPYDNSSYNISP    227
PfGB4_010017100-t41_1-p1         GINKMYKETIHKTLTSDVSTENSYTHNNSRDDEPQNGKRTYNNQSNNNLPYDNSSYNISP    227
SOS76152.1                       GINKMYKETIHKTLTSDVSTENSYTHNNSRDDEPQNGKRTYNNQSNNNLPYDNSSYNISP    227
PfHB3_010017200-t41_1-p1         GINKMYKETIHKTLTSDVSTENSYTHNNSRDDEPQNGKRTYNNQSNNNLPYDNSSYNISP    227
PfTG01_010018100-t41_1-p1        GINKMYKETIHKTLTSDVSTENSYTHNNSRDDEPQNGKRTYNNQSNNNLPYDNSSYNISP    227
PfNF135_010017200.1-p1           GINKMYKETIHKTLTSDVSTENSYTHNNSRDDEPQNGKRTYNNQSNNNLPYDNSSYNISP    227
PfKH02_010016900-t41_1-p1        GINKMYKETIHKTLTSDVSTENSYTHNNSRDDEPQNGKRTYNNQSNNNLPYDNSSYNISP    227
PfNF166_010016800.1-p1           GINKMYKETIHKTLTSDVSTENSYTHNNSRDDEPQNGKRTYNNQSNNNLPYDNSSYNISP    227
Pf7G8-2_000046900.1-p1           GINKMYKETIHKTLTSDVSTENSYTHNNSRDDEPQNGKRTYNNQSNNNLPYDNSSYNISP    227
Pf7G8_010017300-t41_1-p1         GINKMYKETIHKTLTSDVSTENSYTHNNSRDDEPQNGKRTYNNQSNNNLPYDNSSYNISP    227
PfKE01_010016300-t41_1-p1        GINKMYKETIHKTLTSDVSTENSYTHNNSRDDEPQNGKRTYNNQSNNNLPYDNSSYNISP    227
PfIT_010016400-t41_1-p1          GINKMYKETIHKTLTSDVSTENSYTHNNSRDDEPQNGKRTYNNQSNNNLPYDNSSYNISP    227
PfSN01_010015800-t41_1-p1        GINKMYKETIHKTLTSDVSTENSYTHNNSRDDEPQNGKRTYNNQSNNNLPYDNSSYNISP    227
PfCD01_010017400-t41_1-p1        GINKMYKETIHKTLTSDVSTENSYTHNNSRDDEPQNGKRTYNNQSNNNLPYDNSSYNISP    227
PfSD01_010016600-t41_1-p1        GINKMYKETIHKTLTSDVSTENSYTHNNSRDDEPQNGKRTYNNQSNNNLPYDNSSYNISP    227
PfKH01_010018100-t41_1-p1        GINKMYKETIHKTLTSDVSTENSYTHNNSRDDEPQNGKRTYNNQSNNNLPYDNSSYNISP    227
PfGA01_010017200-t41_1-p1        GINKMYKETIHKTLTSDVSTENSYTHNNSRDDEPQNGKRTYNNQSNNNLPYDNSSYNISP    227
PfGN01_010017400-t41_1-p1        GINKMYKETIHKTLTSDVSTENSYTHNNSRDDEPQNGKRTYNNQSNNNLPYDNSSYNISP    227
PfDd2_010016800-t41_1-p1         GINKMYKETIHKTLTSDVSTENSYTHNNSRDDEPQNGKRTYNNQSNNNLPYDNSSYNISP    227
PfML01_010016900-t41_1-p1        GINKMYKETIHKTLTSDVSTENSYTHNNSRDDEPQNGKRTYNNQSNNNLPYDNSSYNISP    227
PRCDC_0111600.1-p1               GINKIYKETIHKALTSDVSTENSNTHNNSRDGEPQNGKLTYNNQSNNNLTYDNSSYNITP    280
PRG01_0113700-t36_1-p1           GINKIYKETIHKALTSDVSTENSNTHNNSRDGEPQNGKLTYNNQSNNNLTYDNSSYNITP    284
                                  :*::::***:::** : **:*: *:**  . : ***:  * * **:** *  * *:* *

PADL01_0110900-t36_1-p1          YN---------DQNNLSQYETVDSNHCNDLNIYNPYYESIDLPYHESTDLPYHEPTDLPY    313
PGABG01_0111700-t36_1-p1         YN---------DQNNLSEYETLDSNHCNDLNIYNPYYQPI--------------------    290
PGSY75_0113300-t31_1-p1          YN---------DQNNLSQYETLDSNHCNDLNIYNPYYQPI--------------------    291
SPJ08214.1                       YNGSNNHVPYNTSNNFSQYNNPDNKYCYYLNIYHPYYGSMNYPHHRLEHYSH---H----    298
PF3D7_0113300.1-p1               YHGPNNNVPYNKSNNFEQCNTQDNKHCNDLDTYHTCYGPDNYPQKYDNYRQECDNYRQEC    287
PfNF54_010017900.1-p1            YHGPNNNVPYNKSNNFEQCNTQDNKHCNDLDTYHTCYGPDNYPQKYDNYRQECDNYRQEC    287
PfGB4_010017100-t41_1-p1         YHGPNNNVPYNKSNNFEQCNTQDNKHCNDLDTYHTCYGPDNYPQKYDNYRQEC-------    280
SOS76152.1                       YHGPNNNVPYNKSNNFEQCNTQDNKHCNDLDTYHTCYGPDNYPQKYDNYRQEYDNYRQEC    287
PfHB3_010017200-t41_1-p1         YHGPNNNVPYNKSNNFEQCNTQDNKHCNDLDTYHTCYGPDNYPQKYDNY-------RQEC    280
PfTG01_010018100-t41_1-p1        YHGPNNNVPYNKSNNFEQCNTQDNKHCNDLDTYHTCYGPDNYPQKYDNY-------RQEC    280
PfNF135_010017200.1-p1           YHGPNNNVPYNKSNNFEQCNTQDNKHCNDLDTYYTCYGPDNYPQKYDNY-------RQEC    280
PfKH02_010016900-t41_1-p1        YHGPNNNVPYNKSNNFEQCNTQDNKHCNDLDTYHTCYGPDNYPQKYDNY-------RQEC    280
PfNF166_010016800.1-p1           YHGPNNNVPYNKSNNFEQCNTQDNKHCNDLDTYHTCYGPDNYPQKYDNY-------RQEC    280
Pf7G8-2_000046900.1-p1           YHGPNNNVPYNKSNNFEQCNTQDNKHCNDLDTYHTCYGPDNYPQKYDNY-------RQEC    280
Pf7G8_010017300-t41_1-p1         YHGPNNNVPYNKSNNFEQCNTQDNKHCNDLDTYHTCYGPDNYPQKYDNY-------RQEC    280
PfKE01_010016300-t41_1-p1        YHGPNNNVPYNKSNNFEQCNTQDNKHCNDLDTYHTCYGPDNYPQKYDNY-------RQEC    280
PfIT_010016400-t41_1-p1          YHGPNNNVPYNKSNNFEQCNTQDNKHCNDLDTYHTCYGPDNYPQKYDNY-------RQEC    280
PfSN01_010015800-t41_1-p1        YHGPNNNVPYNKSNNFEQCNTQDNKHCNDLDTYHTCYGPDNYPQKYDNY-------RQEC    280
PfCD01_010017400-t41_1-p1        YHGPNNNVPYNKSNNFEQCNTQDNKHCNDLDTYHTCYGPDNYPQKYDNY-------RQEC    280
PfSD01_010016600-t41_1-p1        YHGPNNNVPYNKSNNFEQCNTQDNKHCNDLDTYHTCYGPDNYPQKYDNY-------RQEC    280
PfKH01_010018100-t41_1-p1        YHGPNNNVPYNKSNNFEQCNTQDNKHCNDLDTYYTCYGPDNYPQKYDNY-------RQEC    280
PfGA01_010017200-t41_1-p1        YHGPNNNVPYNKSNNFEQCNTQDNKHCNDLDTYHTCYGPDNYPQKYDNY-------RQEC    280
PfGN01_010017400-t41_1-p1        YHGPNNNVPYNKSNNFEQCNTQDNKHCNDLDTYHTCYGPDNYPQKYDNY-------RQEC    280
PfDd2_010016800-t41_1-p1         YHGPNNNVPYNKSNNFEQCNTQDNKHCNDLDTYHTCYGPDNYPQKYDNY-------RQEC    280
PfML01_010016900-t41_1-p1        YHGPNNNVPYNKSNNFEQCNTQDNKHCNDLDTYHTCYGPDNYPQKYDNY-------RQEC    280
PRCDC_0111600.1-p1               YHGPNNNIPYNKSNNSEQCNTQDNKHCNALDIYHTSYGPDNYPHGYDNHTHGYDNHPHGY    340
PRG01_0113700-t36_1-p1           YHGPNNNIPYNKSNNSEQCNTQDNKHCNALDIYHTSYGPDNYPHGYDNHTHGYDNHPHGY    344
                                 *:          .** .: :. *.::*  *: *   *
```

```
PADL01_0110900-t36_1-p1         H----EPTDLPYH---EPTDLP-------YY--EPVSYTHDRSNNYSNRNYSNNN-Y---     353
PGABG01_0111700-t36_1-p1        -------------------DLP-------YY--EPVSYAHDRSNNYSHHSNHNYSHH---     319
PGSY75_0113300-t31_1-p1         -------------------DLP-------YY--EPVSYAHDRSNNYSHHSNHNYSHH---     320
SPJ08214.1                      ---------RPDHYSH-------------------------------HRPDH--------     310
PF3D7_0113300.1-p1              DNYRQEYDNYPQKYDNYRQECDNYRQ----------------------------------     313
PfNF54_010017900.1-p1           DNYRQEYDNYPQKYDNYRQECDNYRQ----------------------------------     313
PfGB4_010017100-t41_1-p1        DNYRQEYDNYPQKYDNYRQECDNYRQ----------------------------------     306
SOS76152.1                      DNYPQEYDNYPQEYDNYPHGFDNYPRGFDNYPRGFDN-----------------------     324
PfHB3_010017200-t41_1-p1        DNYRQEYDNYPHGFDNYPRGFDNYPRGFDNYPHGFDN-----------------------     317
PfTG01_010018100-t41_1-p1       DNYRQECDNYRQEYDNYPHGFDNYPRGFDNYPHGFDN-----------------------     317
PfNF135_010017200.1-p1          DNYRQKYDNYPHGFDNYPRGFDNYPHGFDNYPRGF-------------------------     315
PfKH02_010016900-t41_1-p1       DNYRQECDNYPHGFDNYPRGFDNYPRGFDNYPRGF-------------------------     315
PfNF166_010016800.1-p1          DNYRQEYDNYPHGFDNYPRGFDNYPHGFDNYPRGF-------------------------     315
Pf7G8-2_000046900.1-p1          DNYRQEYDNYPHGFDNYPRGFDNYPRGFDNYPRGFDN-----------------------     317
Pf7G8_010017300-t41_1-p1        DNYRQEYDNYPHGFDNYPRGFDNYPRGFDNYPRGFDN-----------------------     317
PfKE01_010016300-t41_1-p1       DNYRQECDNYPHGFDNYPRGFDNYPRGFDNYPRGFDN-----------------------     317
PfIT_010016400-t41_1-p1         DNYRQEYDNYPHGFDNYPHGFDNYP-------RGFDN-----------------------     310
PfSN01_010015800-t41_1-p1       DNYRQEYDNYPHGFDNYPRGFDNYP-------RGFDN-----------------------     310
PfCD01_010017400-t41_1-p1       DNYRQEYDNYPHGFDNYPRGFDNYP-------RGFDN-----------------------     310
PfSD01_010016600-t41_1-p1       DNYRQEYDNYPHGFDNYPRGFDNYP-------RGFDN-----------------------     310
PfKH01_010018100-t41_1-p1       DNYRQEYDNYPHGFDNYPRGFDNYP-------R---------------------------     306
PfGA01_010017200-t41_1-p1       DNYRKECDNYPHGFDNYPRGFDNYP-------R---------------------------     306
PfGN01_010017400-t41_1-p1       DNYRQECDNYPHGFDNYPRGFDNYP-------R---------------------------     306
PfDd2_010016800-t41_1-p1        DNYRQEYDNYPHGFDNYPRGFDNYP-------H---------------------------     306
PfML01_010016900-t41_1-p1       DNYRQEYDNYPHGFDNYPRGFDNYP-------H---------------------------     306
PRCDC_0111600.1-p1              DNHTHGYDNHPHGYDNHPHGYDKHPHGPDNHP----------------------------     372
PRG01_0113700-t36_1-p1          DNHPHGYDNHPHGYDNHPHGYDKHPHGYDNYPHRPHIYPHG-YDNYPHRPHIYPHGYDNH     403


PADL01_0110900-t36_1-p1         --SNRNY------SNNNYSKHNYPHH-------------NYSHHNYPHHNYPHHNYPHHN     392
PGABG01_0111700-t36_1-p1        --SNHNYHHHSNHNHSHHSNHNYPHH-------------SNHNHNHSHHNHSH-----HN     359
PGSY75_0113300-t31_1-p1         --SNHNYSHHSNHNYSHHSNHNYPHH-------------SNHNHNHSHHNHPHHNHSHHN     365
SPJ08214.1                      ------YSH--------HKPDHYLH------HKPDHYL------HHKPDHY------LHH     338
PF3D7_0113300.1-p1              ------------------EYDNYPHGFDNYPRGFDNYPHGYDNHPHRPHIYPHGFDNHPH     355
PfNF54_010017900.1-p1           ------------------EYDNYPHGFDNYPRGFDNYPHGYDNHPHRPHIYPHGFDNHPH     355
PfGB4_010017100-t41_1-p1        ------------------EYDNYPHGFDNYPRGFDNYPHGYDNHPHRPHIYPHGFDNHPH     348
SOS76152.1                      ------YPRGF--DNYPHGFDNYPHGFDNYPHGFDNYPHGYDNHPHRPHIYPHGFDNHPH     376
PfHB3_010017200-t41_1-p1        ------YPHGF--DNYPHGFDNYPHGYDNHPHRPHIYPHGFDNHPHRPHIYPHGFDNHPH     369
PfTG01_010018100-t41_1-p1       ------YPHGF--DNYPHGFDNYPHGYDNHPHRPHIY---------------PHGFDNHPH     355
PfNF135_010017200.1-p1          -------------DNYPRGFDNYPHGYDNHPHRPHIY---------------PHGFDNHPH     348
PfKH02_010016900-t41_1-p1       -------------DNYPRGFDNYPHGYDNHPHRPHIY---------------PHGFDNHPH     348
PfNF166_010016800.1-p1          -------------DNYPRGFDNYPHGYDNHPHRPHIY---------------PHGFDNHPH     348
Pf7G8-2_000046900.1-p1          ------YPHGF--DNYPHGFDNYPHGYDNHPHRPHIY---------------PHGFDNHPH     355
Pf7G8_010017300-t41_1-p1        ------YPHGF--DNYPHGFDNYPHGYDNHPHRPHIY---------------PHGFDNHPH     355
PfKE01_010016300-t41_1-p1       ------YPHGF--DNYPHGFDNYPHGYDNHPHRPHIY---------------PHGFDNHPH     355
PfIT_010016400-t41_1-p1         ------YPHGF--DNYPHGFDNYPHGYDNHPHRPHIY---------------PHGFDNHPH     348
PfSN01_010015800-t41_1-p1       ------YPHGF--DNYPHGFDNYPHGYDNHPHRPHIY---------------PHGFDNHPH     348
PfCD01_010017400-t41_1-p1       ------YPHGF--DNYPHGFDNYPHGYDNHPHRPHIY---------------PHGFDNHPH     348
PfSD01_010016600-t41_1-p1       ------YPHGF--DNYPHGFDNYPHGYDNHPHRPHIY---------------PHGFDNHPH     348
PfKH01_010018100-t41_1-p1       ---------GF--DNYPRGFDNYPHGYDNHPHRPHIY---------------PHGFDNHPH     341
PfGA01_010017200-t41_1-p1       ---------GF--DNYPRGFDNYPHGYDNHPHRPHIY---------------PHGFDNHPH     341
PfGN01_010017400-t41_1-p1       ---------GF--DNYPHGFDNYPHGYDNHPHRPHIY---------------PHGFDNHPH     341
PfDd2_010016800-t41_1-p1        ---------GF--DNYPHGFDNYPHGYDNHPHRPHIY---------------PHGFDNHPH     341
PfML01_010016900-t41_1-p1       ---------GF--DNYPHGFDNYPHGYDNHPHRPHIY---------------PHGFDNHPH     341
PRCDC_0111600.1-p1              --------HGY--DNHPHGYDNHPHGYDNYPHRPHIYPHGYDNHPHRPHIYPHGYDNHPH     422
PRG01_0113700-t36_1-p1          PHRPHIYPHGY--DNHPHRPHIYPHGYDNHPHRPHIYPHGYDNHPHRPHIYPHGYDNHPH     461
                                                    . : *                                  :
```

```
PADL01_0110900-t36_1-p1         YPHH---NYPHHRYGSSGDPYHRHDHIMDRSYYYSNLQNDNHDMMLTYIPKNDNKSLYDE    449
PGABG01_0111700-t36_1-p1        HSHHNHSHRSHHRYGPGGDPYHRHDHIMDRSYYNNNIQNDNDDMMLTYNQMNDNKSLYDE    419
PGSY75_0113300-t31_1-p1         HSHHNHSHHNHHRYGPGGDPYHRHDHIMDRSYYNNNLQNDNDDMMLTYNQMNDNKSLYDE    425
SPJ08214.1                      RQ-NKYSHNLSLRNISVGGPYYRPAHMMERFDYYSNLKNDAHYMMLPYNRMNDNKSMCDE    397
PF3D7_0113300.1-p1              RP-HMYPHNFPMRNESVGGPYYRPPHIIERSNYYKNPKKAPHNMMLPCDTMKDNKSICDE    414
PfNF54_010017900.1-p1           RP-HMYPHNFPMRNESVGGPYYRPPHIIERSNYYKNPKKAPHNMMLPCDTMKDNKSICDE    414
PfGB4_010017100-t41_1-p1        RP-HMYPHNFPMRNESVGGPYYRPPHIIERSNYYRNPKKAPHNMMLPCDTMKDNKSICDE    407
SOS76152.1                      RP-HMYPHNFPMRNESVGGPYYRPPHIIERSNYYRNPKKAPHNMMLPCDTMKDNKSICDE    435
PfHB3_010017200-t41_1-p1        RP-HMYPHNFPMRNESVGGPYYRPPHIIERSNYYRNPKKAPHNMMLPCDTMKDNKSICDE    428
PfTG01_010018100-t41_1-p1       RP-HMYPHNFPMRNESVGGPYYRPPHIIERSNYYRNPKKAPHNMMLPCDTMKDNKSICDE    414
PfNF135_010017200.1-p1          RP-HMYPHNFPMRNESVGGPYYRPPHIIERSNYYRNPKKAPHNMMLPCDTMKDNKSICDE    407
PfKH02_010016900-t41_1-p1       RP-HMYPHNFPMRNESVGGPYYRPPHIIERSNYYRNPKKAPHNMMLPCDTMKDNKSICDE    407
PfNF166_010016800.1-p1          RP-HMYPHNFPMRNESVGGPYYRPPHIIERSNYYRNPKKAPHNMMLPCDTMKDNKSICDE    407
Pf7G8-2_000046900.1-p1          RP-HMYPHNFPMRNESVGGPYYRPPHIIERSNYYKNPKKAPHNMMLPCDTMKDNKSICDE    414
Pf7G8_010017300-t41_1-p1        RP-HMYPHNFPMRNESVGGPYYRPPHIIERSNYYKNPKKAPHNMMLPCDTMKDNKSICDE    414
PfKE01_010016300-t41_1-p1       RP-HMYPHNFPMRNESVGGPYYRPPHIIERSNYYRNPKKAPHNMMLPCDTMKDNKSICDE    414
PfIT_010016400-t41_1-p1         RP-HMYPHNFPMRNESVGGPYYRPPHIIERSNYYRNPKKAPHNMMLPCDTMKDNKSICDE    407
PfSN01_010015800-t41_1-p1       RP-HMYPHNFPMRNESVGGPYYRPPHIIERSNYYRNPKKAPHNMMLPCDTMKDNKSICDE    407
PfCD01_010017400-t41_1-p1       RP-HMYPHNFPMRNESVGGPYYRPPHIIERSNYYRNPKKAPHNMMLPCDTMKDNKSICDE    407
PfSD01_010016600-t41_1-p1       RP-HMYPHNFPMRNESVGGPYYRPPHIIERSNYYRNPKKAPHNMMLPCDTMKDNKSICDE    407
PfKH01_010018100-t41_1-p1       RP-HMYPHNFPMRNESVGGPYYRPPHIIERSNYYRNPKKAPHNMMLPCDTMKDNKSICDE    400
PfGA01_010017200-t41_1-p1       RP-HMYPHNFPMRNESVGGPYYRPPHIIERSNYYRNPKKAPHNMMLPCDTMKDNKSICDE    400
PfGN01_010017400-t41_1-p1       RP-HMYPHNFPMRNESVGGPYYRPPHIIERSNYYRNPKKAPHNMMLPCDTMKDNKSICDE    400
PfDd2_010016800-t41_1-p1        RP-HMYPHNFPMRNESVGGPYYRPPHIIERSNYYRNPKKAPHNMMLPCDTMKDNKSICDE    400
PfML01_010016900-t41_1-p1       RP-HMYPHNFPMRNESVGGPYYRPPHIIERSNYYRNPKKAPHNMMLPCDTMKDNKSICDE    400
PRCDC_0111600.1-p1              RP-HIYPHNFPMRNESVGGPYYRPPHLIERSDYYSNPKNEPHNMMLPYNTMEDNTSICDE    481
PRG01_0113700-t36_1-p1          RP-HIYPHNFPMRNESVGGPYYRPPHLIERSDYYSNPKNEPHNMMLPYNTMEDNTSICDE    520
                                   :   :    *    *.**:*  *:::*  *  * ::  . ***     :**.*: **

PADL01_0110900-t36_1-p1         QNFELELKKIIKKNHLQNDNITDSYDTAVNDFNKKLKEYNKKLNEYNEKLNEYTS-RLNE    508
PGABG01_0111700-t36_1-p1        QNFELELKKIIKKNHLQNSNITDSCDTRVSDYNKKLVEYNKKLNEYNEKLNEYTR-RLNE    478
PGSY75_0113300-t31_1-p1         QNFELELKKIIKKNHLQNGNITDSCDTRVSDYNKKLVEYNKKLNEYNEKLNEYTR-RLNE    484
SPJ08214.1                      QNFQLGLEKMKKKNNLQNGNIRDSHDTRISDYNKRLNGYNKRLNGYNKKNSYNKQSSYNK    457
PF3D7_0113300.1-p1              QNFQRELEKIIKKNNLQNGNIRDNHDTRINDYNKRLTEYNKRLTEYNKRLTEYTK-RLNE    473
PfNF54_010017900.1-p1           QNFQRELEKIIKKNNLQNGNIRDNHDTRINDYNKRLTEYNKRLTEYNKRLTEYTK-RLNE    473
PfGB4_010017100-t41_1-p1        QNFQRELEKIIKKNNLQNGNIRDNHDTRINDYNKRLTEYNKRLTEYNKRLTEYTK-RLNE    466
SOS76152.1                      QNFQRELEKIIKKNNLQNGNSRDNHDTRINDYNKRLTEYNKRLTEYNKRLTEYTK-RLNE    494
PfHB3_010017200-t41_1-p1        QNFQRELEKIIKKNNLQNGNIRDNHDTRINDYNKRLTEYNKRLTAYNKRLTEYTK-RLNE    487
PfTG01_010018100-t41_1-p1       QNFQRELEKIIKKNNLQNGNIRDNHDTRINDYNKRLTEYNKRLTEYNKRLTEYTK-RLNE    473
PfNF135_010017200.1-p1          QNFQRELEKIIKKNNLQNGNIRDNHDTRINDYNKRLTEYNKRLTEYNKRLTEYTK-RLNE    466
PfKH02_010016900-t41_1-p1       QNFQRELEKIIKKNNLQNGNIRDNHDTRINDYNKRLTEYNKRLTEYNKRLTEYTK-RLNE    466
PfNF166_010016800.1-p1          QNFQRELEKIIKKNNLQNGNIRDNHDTRINDYNKRLTEYNKRLTEYNKRLTEYTK-RLNE    466
Pf7G8-2_000046900.1-p1          QNFQRELEKIIKKNNLQNGNIRDNHDTRINDYNKRLTEYNKRLTEYNKRLTEYTK-RLNE    473
Pf7G8_010017300-t41_1-p1        QNFQRELEKIIKKNNLQNGNIRDNHDTRINDYNKRLTEYNKRLTEYNKRLTEYTK-RLNE    473
PfKE01_010016300-t41_1-p1       QNFQRELEKIIKKNNLQNGNIRDNHDTRINDYNKRLTEYNKRLTEYNKRLTEYTK-RLNE    473
PfIT_010016400-t41_1-p1         QNFQRELEKIIKKNNLQNGNIRDNHDTRINDYNKRLTEYNKRLTEYNKRLTEYTK-RLNE    466
PfSN01_010015800-t41_1-p1       QNFQRELEKIIKKNNLQNGNIRDNHDTRINDYNKRLTEYNKRLTEYNKRLTEYTK-RLNE    466
PfCD01_010017400-t41_1-p1       QNFQRELEKIIKKNNLQNGNIRDNHDTRINDYNKRLTEYNKRLTEYNKRLTEYTK-RLNE    466
PfSD01_010016600-t41_1-p1       QNFQRELEKIIKKNNLQNGNIRDNHDTRINDYNKRLTEYNKRLTEYNKRLTEYTK-RLNE    466
PfKH01_010018100-t41_1-p1       QNFQRELEKIIKKNNLQNGNIRDNHDTRINDYNKRLTEYNKRLTEYNKRLTEYTK-RLNE    459
PfGA01_010017200-t41_1-p1       QNFQRELEKIIKKNNLQNGNIRDNHDTRINDYNKRLTEYNKRLTEYNKRLTEYTK-RLNE    459
PfGN01_010017400-t41_1-p1       QNFQRELEKIIKKNNLQNGNIRDNHDTRINDYNKRLTEYNKRLTEYNKRLTEYTK-RLNE    459
PfDd2_010016800-t41_1-p1        QNFQRELEKIIKKNNLQNGNIRDNHDTRINDYNKRLTEYNKRLTEYNKRLTEYTK-RLNE    459
PfML01_010016900-t41_1-p1       QNFQRELEKIIKKNNLQNGNIRDNHDTRINDYNKRLTEYNKRLTEYNKRLTEYTK-RLNE    459
PRCDC_0111600.1-p1              QSFQLELEKIIKKNNLQNGNIRDSHDTGINDYDKRLTEYNKRLTEYNKRLNEYTK-RLNE    540
PRG01_0113700-t36_1-p1          QSFQLELEKIIKKNNLQNGNIRDSHDTGINDYDKRLTEYNKRLTEYNKRLNEYTK-RLNE    579
                                *.*:  *:*: ***:***.*  *. ** :.*::*:*  ***:*. **:: .  .    *:
```

```
PADL01_0110900-t36_1-p1       YNKKHNEKKKNDDNKKSGQNNNENILSQDIVLYGTDFQNAFRYRQNTRSYYSNIYSNREE    568
PGABG01_0111700-t36_1-p1      YNKKHNEN------KQSRQNNNENTLSQDIVLYGADFQNAFRYRQNTRSYYSNIHSNREE    532
PGSY75_0113300-t31_1-p1       YNKKHNEN------KQSRQNNNENTLSQDIVLYGADFQNAFRYRQNTRSYYSNIHSNREE    538
SPJ08214.1                    QSS------YNKQNCYNKQNGIENRSSNNIVLYGNNFQNAFRYRQNTRSYYTNIYSNGEA    511
PF3D7_0113300.1-p1            HYK------RNGYNIQNRQNSIERAQSNDVVLYGHNFQNAFRYKQNTRSYYPHVNSNEAT    527
PfNF54_010017900.1-p1         HYK------RNGYNIQNRQNSIERAQSNDVVLYGHNFQNAFRYKQNTRSYYPHVNSNEAT    527
PfGB4_010017100-t41_1-p1      HYK------RKGYNIQNRQNSIERAQSNDVVLYGHNFQNAFRYKQNTRSYYPHVNSNEAT    520
SOS76152.1                    HYK------RNGYNIQNRQNSIERAPSNDVVLYGHNFQNAFRYKQNTRSYYPHVNSNEAT    548
PfHB3_010017200-t41_1-p1      HYK------RNGYNIQNRQNSIERAQSNDVVLYGHNFQNAFRYKQNTRSYYPHVNSNEAT    541
PfTG01_010018100-t41_1-p1     HYK------RKGYNIQNRQNSIERAQSNDVVLYGHNFQNAFRYKQNTRSYYPHVNSNEAT    527
PfNF135_010017200.1-p1        HYK------RNGYNIQNRQNSIERAQSNDVVLYGHNFQNAFRYKQNTRSYYPHVNSNEAT    520
PfKH02_010016900-t41_1-p1     HYK------RNGYNIQNRQNSIERAQSNDVVLYGHNFQNAFRYKQNTRSYYPHVNSNEAT    520
PfNF166_010016800.1-p1        HYK------RKGYNIQNRQNSIERAQSNDVVLYGHNFQNAFRYKQNTRSYYPHVNSNEAT    520
Pf7G8-2_000046900.1-p1        HYK------RKGYNIQNRQNSIERAQSNDVVLYGHNFQNAFRYKQNTRSYYPHVNSNEAT    527
Pf7G8_010017300-t41_1-p1      HYK------RKGYNIQNRQNSIERAQSNDVVLYGHNFQNAFRYKQNTRSYYPHVNSNEAT    527
PfKE01_010016300-t41_1-p1     HYK------RKGYNIQNRQNSIERAQSNDVVLYGHNFQNAFRYKQNTRSYYPHVNSNEAT    527
PfIT_010016400-t41_1-p1       HYK------RNGYNIQNRQNSIERAQSNDVVLYGHNFQNAFRYKQNTRSYYPHVNSNEAT    520
PfSN01_010015800-t41_1-p1     HYK------RNGYNIQNRQNSIERAQSKDVVLYGHNFQNAFRYKQNTRSYYPHVNSNEAT    520
PfCD01_010017400-t41_1-p1     HYK------RNGYNIQNRQNSIERAQSKDVVLYGHNFQNAFRYKQNTRSYYPHVNSNEAT    520
PfSD01_010016600-t41_1-p1     HYK------RNGYNIQNRQNSIERAQSNDVVLYGHNFQNAFRYKQNTRSYYPHVNSNEAT    520
PfKH01_010018100-t41_1-p1     HYK------RNGYNIQNRQNSIERAQSNDVVLYGHNFQNAFRYKQNTRSYYPHVNSNEAT    513
PfGA01_010017200-t41_1-p1     HYK------RKGYNIQNRQNSIERAQSNDVVLYGHNFQNAFRYKQNTRSYYPHVNSNEAT    513
PfGN01_010017400-t41_1-p1     HYK------RNGYNIQNRQNSIERAQSNDVVLYGHNFQNAFRYKQNTRSYYPHVNSNEAT    513
PfDd2_010016800-t41_1-p1      HYK------RNGYNIQNRQNSIERAQSNDVVLYGHNFQNAFRYKQNTRSYYPHVNSNEAT    513
PfML01_010016900-t41_1-p1     HYK------RKGYNIQNRQNSIERAQSNDVVLYGHNFQNAFRYKQNTRSYYPHVNSNEAT    513
PRCDC_0111600.1-p1            HYK------RSGYNIQNRQNSIEHIPSNDIVLYGHNFQNAFRYKQNTRSYYPHVNSKGTA    594
PRG01_0113700-t36_1-p1        HYK------RSGYNIQNRQNSIEHIPSNDIVLYGHNFQNAFRYKQNTRSYYPHVNSKGTA    633
                                .             . **. *.  *:::**** :*******:******* :: *:

PADL01_0110900-t36_1-p1       DLDKVIDFTHNNNSSEDEYTFRNKQEIYHQHSKRLEKKLFDYQNGSNPVINFLERHF       625
PGABG01_0111700-t36_1-p1      DLDRVIDFTNNNNNSSEDEYTFGNEQEIYHQHSKRLEKKLFDYQNGTNPLINFLERHF      589
PGSY75_0113300-t31_1-p1       DLDRVIDFTNNNNNASEDEYTFGNEQEIYHQHSKRLEKKLFDYQNGTNPLINFLERHF      595
SPJ08214.1                    NHERARYFNQQNNRPQEEYPIKSEQHMHHVQSKRLDKKLYDYENCSNPVRNFLKRHF       568
PF3D7_0113300.1-p1            HHQKTMYFTQQNNYSREEYPIKSEQHLYHVKSKRLEKKLYDYQNGTNPVTNFLERHF       584
PfNF54_010017900.1-p1         HHQKTMYFTQQNNYSREEYPIKSEQHLYHVKSKRLEKKLYDYQNGTNPVTNFLERHF       584
PfGB4_010017100-t41_1-p1      HHQKTMYFTQQNNYSREEYPIKSEQHLYHVKSKRLEKKLYDYQNGTNPVTNFLERHF       577
SOS76152.1                    HHQKTMYFTQQNNYSREEYPIKSEQHLYHVKSKRLEKKLYDYQNGTNPVTNFLERHF       605
PfHB3_010017200-t41_1-p1      HHQKTMYFTQQNNYSREEYPIKSEQHLYHVKSKRLEKKLYDYQNGTNPVTNFLERHF       598
PfTG01_010018100-t41_1-p1     HHQKTMYFTQQNNYSREEYPIKSEQHLYHVKSKRLEKKLYDYQNGTNPVTNFLERHF       584
PfNF135_010017200.1-p1        HHQKTMYFTQQNNYSREEYPIKSEQHLYHVKSKRLEKKLYDYQNGTNPVTNFLERHF       577
PfKH02_010016900-t41_1-p1     HHQKTMYFTQQNNYSREEYPIKSEQHLYHVKSKRLEKKLYDYQNGTNPVTNFLERHF       577
PfNF166_010016800.1-p1        HHQKTMYFTQQNNYSREEYPIKSEQHLYHVKSKRLEKKLYDYQNGTNPVTNFLERHF       577
Pf7G8-2_000046900.1-p1        HHQKTMYFTQQNNYSREEYPIKSEQHLYHVKSKRLEKKLYDYQNGTNPVTNFLERHF       584
Pf7G8_010017300-t41_1-p1      HHQKTMYFTQQNNYSREEYPIKSEQHLYHVKSKRLEKKLYDYQNGTNPVTNFLERHF       584
PfKE01_010016300-t41_1-p1     HHQKTMYFTQQNNYSREEYPIKSEQHLYHVKSKRLEKKLYDYQNGTNPVTNFLERHF       584
PfIT_010016400-t41_1-p1       HHQKTMYFTQQNNYSREEYPIKSEQHLYHVKSKRLEKKLYDYQNGTNPVTNFLERHF       577
PfSN01_010015800-t41_1-p1     HHQKTMYFTQQNNYSREEYPIKSEQHLYHVKSKRLEKKLYDYQNGTNPVTNFLERHF       577
PfCD01_010017400-t41_1-p1     HHQKTMYFTQQNNYSREEYPIKSEQHLYHVKSKRLEKKLYDYQNGTNPVTNFLERHF       577
PfSD01_010016600-t41_1-p1     HHQKTMYFTQQNNYSREEYPIKSEQHLYHVKSKRLEKKLYDYQNGTNPVTNFLERHF       577
PfKH01_010018100-t41_1-p1     HHQKTMYFTQQNNYSREEYPIKSEQHLYHVKSKRLEKKLYDYQNGTNPVTNFLERHF       570
PfGA01_010017200-t41_1-p1     HHQKTMYFTQQNNYSREEYPIKSEQHLYHVKSKRLEKKLYDYQNGTNPVTNFLERHF       570
PfGN01_010017400-t41_1-p1     HHQKTMYFTQQNNYSREEYPIKSEQHLYHVKSKRLEKKLYDYQNGTNPVTNFLERHF       570
PfDd2_010016800-t41_1-p1      HHQKTMYFTQQNNYSREEYPIKSEQHLYHVKSKRLEKKLYDYQNGTNPVTNFLERHF       570
PfML01_010016900-t41_1-p1     HHQKTMYFTQQNNYSREEYPIKSEQHLYHVKSKRLEKKLYDYQNGTNPVTNFLERHF       570
PRCDC_0111600.1-p1            QHQKSMYFTQQNNYSQ-EYPTKSEQHMYHVKSKRLEKKLYDYQNGTNPITNFLERHF       650
PRG01_0113700-t36_1-p1        QHQKSMYFTQQNNYSQ-EYPTKSEQHMYHVKSKRLEKKLYDYQNGTNPITNFLERHF       689
                              . ::   *.::**  . **   .:*.::* :****:***:**:* :**: ***:***
```
