## Supplementary figures and images for "A systematic targeted genetic screen identifies proteins involved in cytoadherence of the malaria parasite *P. falciparum*"

### Supplementary 2

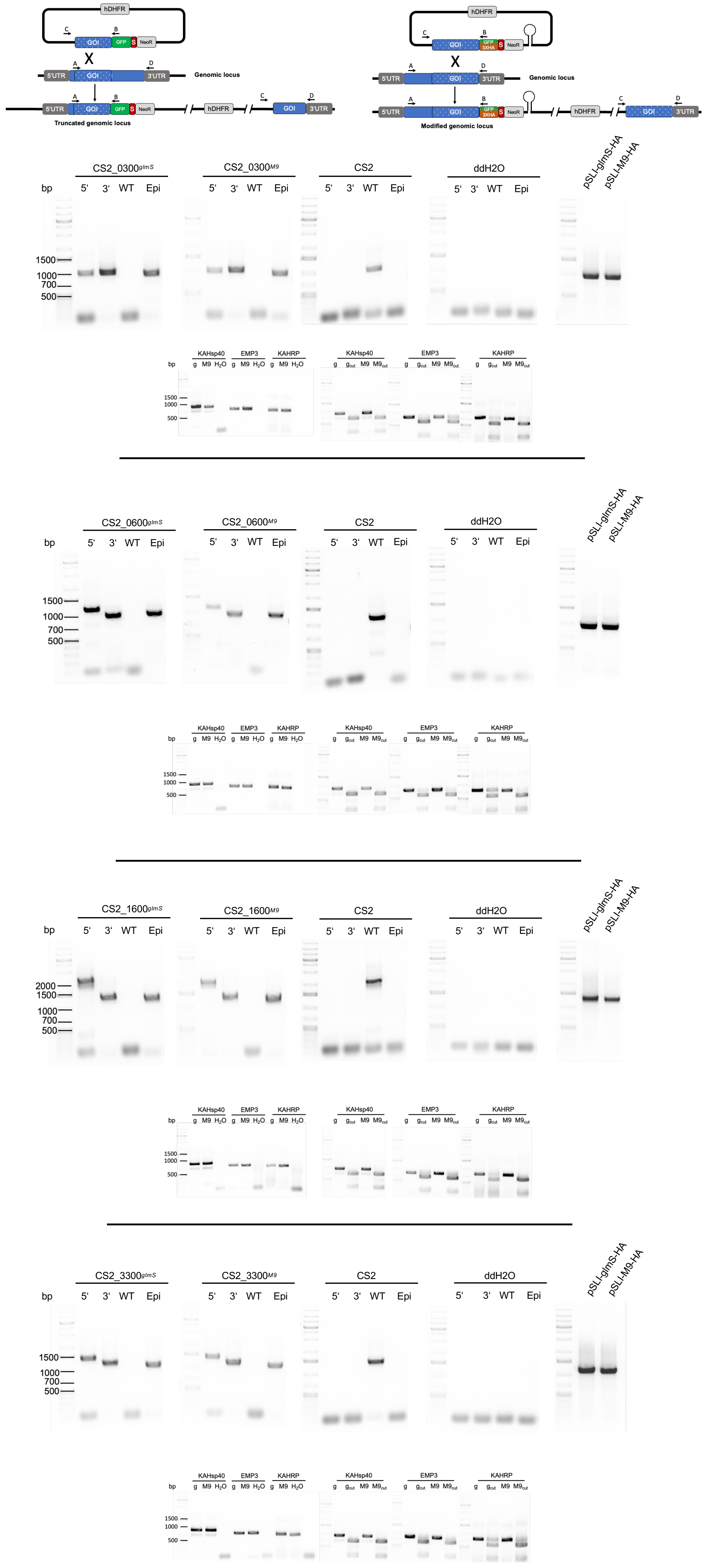

### Supplementary 4

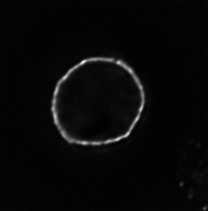

### Supplementary 6

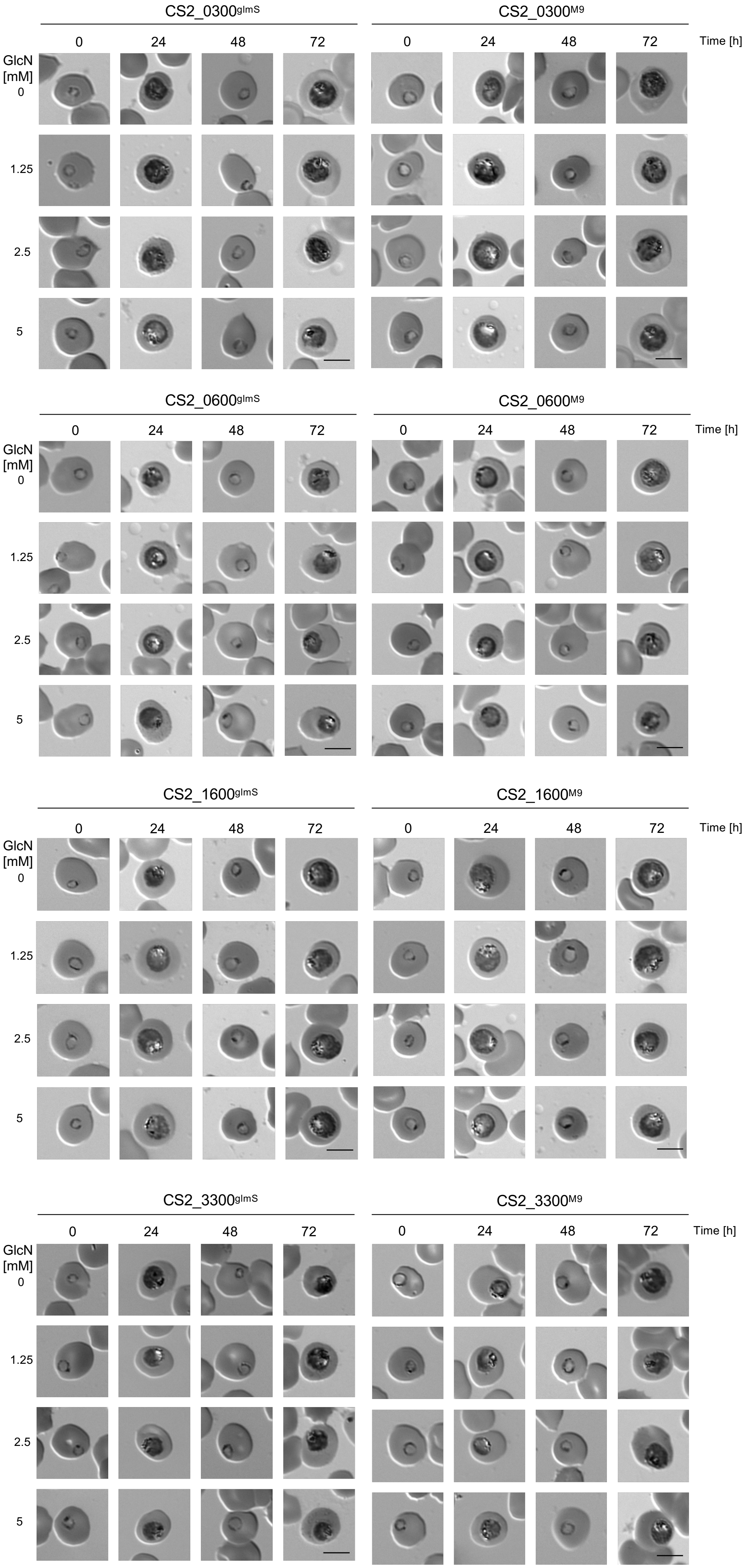

### Supplementary 7

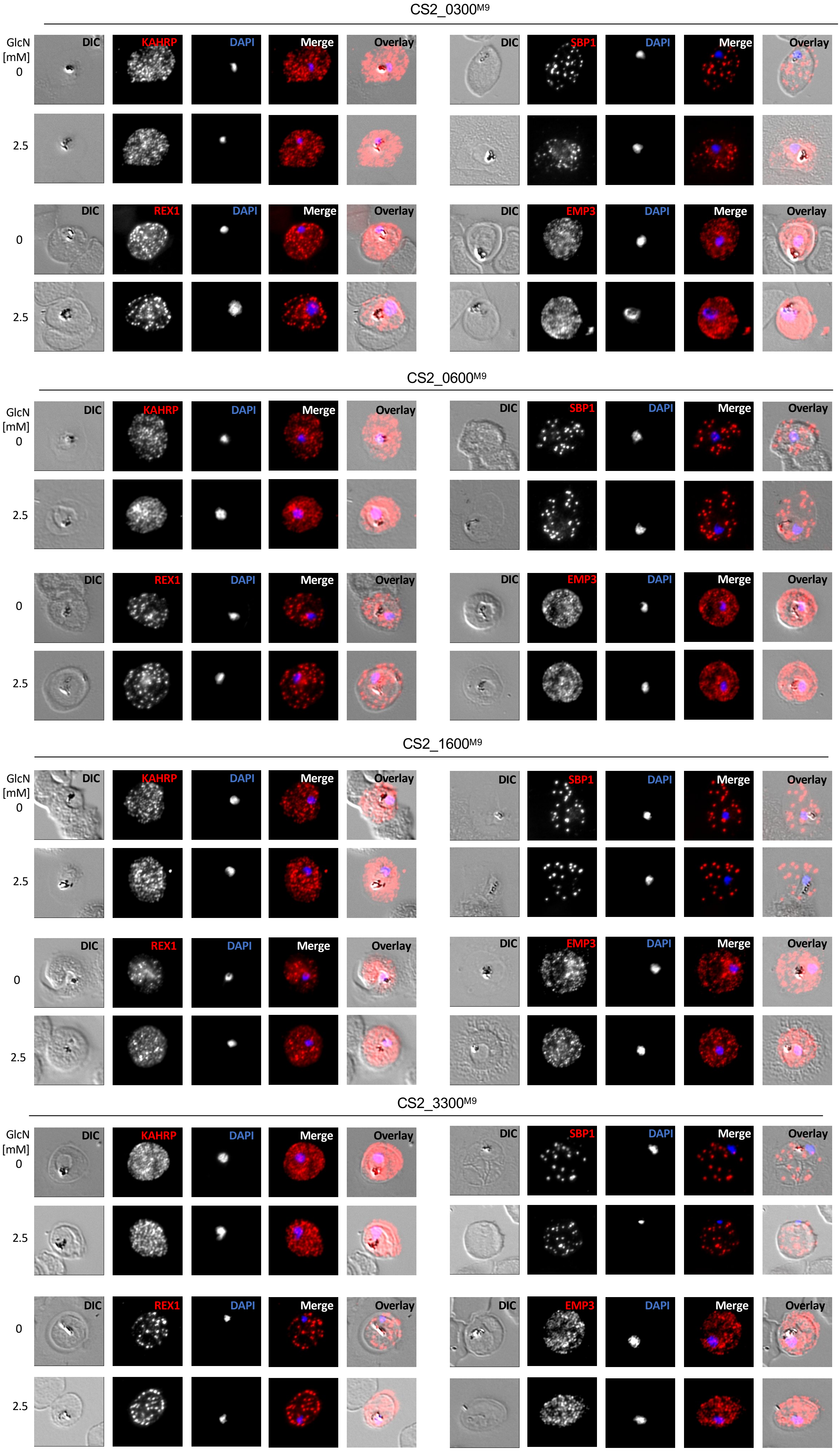
